# Functional impact of the H2A.Z histone variant during meiosis in *Saccharomyces cerevisiae*

**DOI:** 10.1101/316133

**Authors:** Sara González-Arranz, Santiago Cavero, Macarena Morillo-Huesca, Eloisa Andújar, Mónica Pérez-Alegre, Félix Prado, Pedro San-Segundo

**Author notes:** Corresponding author: Pedro A. San Segundo, Instituto de Biología Funcional y Genómica, CSIC-University of Salamanca, Zacarías González, 2, 37007 Salamanca, Spain.

## Abstract

Among the collection of chromatin modifications that influence its function and structure, the substitution of canonical histones by the so-called histone variants is one of the most prominent actions. Since crucial meiotic transactions are modulated by chromatin, here we investigate the functional contribution of the H2A.Z histone variant during both unperturbed meiosis and upon challenging conditions where the meiotic recombination checkpoint is triggered in budding yeast by the absence of the synaptonemal complex component Zip1. We have found that H2A.Z localizes to meiotic chromosomes in an SWR1-dependent manner. Although meiotic recombination is not substantially altered, the *htz1* mutant (lacking H2A.Z) shows inefficient meiotic progression, impaired sporulation and reduced spore viability. These phenotypes are likely accounted for by the misregulation of meiotic gene expression landscape observed in *htz1*. In the *zip1* mutant, the absence of H2A.Z results in a tighter meiotic arrest imposed by the meiotic recombination checkpoint. We have found that Mec1-dependent Hop1-T318 phosphorylation and the ensuing Mek1 activation are not significantly altered in *zip1 htz1*; however, downstream checkpoint targets, such as the meiosis I-promoting factors Ndt80, Cdc5 and Clb1, are drastically down-regulated. The study of the checkpoint response in *zip1 htz1* has also allowed us to reveal the existence of an additional function of the Swe1 kinase, independent of CDK inhibitory phosphorylation, which is relevant to restrain meiotic cell cycle progression. In summary, our study shows that the H2A.Z histone variant impacts various aspects of meiotic development adding further insight into the relevance of chromatin dynamics for accurate gametogenesis.

## Introduction

Sexual reproduction relies on a specialized cell division, meiosis, which reduces chromosome ploidy by half and is usually accompanied by cell differentiation processes that culminate in the formation of gametes. The reduction in chromosome complement is achieved by two consecutive rounds of nuclear division preceded by a single round of DNA replication. Premeiotic S phase is followed by a long prophase I in which, prior to the first meiotic division, homologous chromosomes (homologs) pair, synapse and recombine. Meiotic recombination is initiated by programmed DNA double-strand breaks (DSBs) generated by the Spo11 protein and a cohort of regulatory factors (Keeney *et al.* 2014). During the repair of a subset of these meiotic DSBs, crossovers between homologs are formed, which are essential for correct distribution of chromosomes to the meiotic progeny. Alignment of homologous chromosomes-pairing- and the stabilization of these interactions by the synaptonemal complex (SC)-synapsis-influence meiotic recombination outcomes (Hunter 2015). These crucial meiotic events are monitored by the so-called meiotic recombination checkpoint (MRC), an evolutionarily-conserved surveillance mechanism that senses defective synapsis and/or recombination and imposes a block or delay in meiotic cell progression providing time to fix the faulty process in order to prevent aberrant chromosome segregation. The meiotic checkpoint network also operates in unperturbed meiosis to ensure the proper sequential execution of events (MacQueen and Hochwagen 2011; Subramanian and Hochwagen 2014).

In this work, we have used the *zip1* mutant of the budding yeast *Saccharomyces cerevisiae* as a genetic tool to activate the MRC. Zip1 is a major structural component of the SC central region and *ZIP1* deletion impairs synapsis and crossover recombination (Dong and Roeder 2000; Borner *et al.* 2004; Voelkel-Meiman *et al.* 2015); as a consequence, the *zip1* mutant experiences a significant MRC-dependent delay in the prophase to meiosis I transition (Herruzo *et al.* 2016). The *zip1*-induced defects are detected by the Mec1-Ddc2^(ATR-ATRIP)^ complex resulting in phosphorylation of the Hop1 checkpoint adaptor at several residues, including T318 (Carballo *et al.* 2008; Refolio *et al.* 2011; Penedos *et al.* 2015). The Hop1 protein is a component of the lateral elements of the SC; its abundance, dynamics and phosphorylation state at chromosome axes in response to checkpoint activation are finely tuned by the AAA+ ATPase Pch2 (Herruzo *et al.* 2016). Phosphorylated Hop1 recruits the meiosis-specific Mek1 protein to chromosomes facilitating the activation of this Rad53/Chk2-related kinase containing an FHA domain in two steps: first by Mec1-dependent phosphorylation and subsequently by *in trans* autophosphorylation of Mek1 dimers on its activation loop (Niu *et al.* 2005; Ontoso *et al.* 2013). In turn, active Mek1 stabilizes Hop1-T318 phosphorylation at chromosomes (Chuang *et al.* 2012). Mek1 promotes interhomolog recombination bias by the direct phosphorylation of the recombination mediator Rad54 at T154 to attenuate its interaction with the strand-exchange Rad51 protein (Niu *et al.* 2009). Also, the phosphorylation of Hed1 at Thr40 stabilizes this protein stimulating its inhibitory action on Rad51 (Callender *et al.* 2016). Mek1 also exerts a spatial control on recombination bias by a synapsis-dependent mechanism involving Pch2 (Subramanian *et al.* 2016). In addition, Mek1 is essential for the meiotic checkpoint response to the accumulation of unrepaired DSBs and to the *zip1*-induced synapsis and/or recombination defects (Xu *et al.* 1997; Ontoso *et al.* 2013; Prugar *et al.* 2017). The arrest or delay at meiotic prophase I imposed by the MRC is established by two interconnected mechanisms: down-regulation of the Ndt80 transcription factor and inhibitory phosphorylation of Cdc28^CDK1^ (Subramanian and Hochwagen 2014). Ndt80 is a master regulator of yeast meiotic development that activates the transcription of a number of genes involved in meiotic divisions and spore formation (Winter 2012). Among the gene products regulated by Ndt80, the polo-like kinase Cdc5 and the type-B Clb1 cyclin are crucial factors to promote exit from prophase (Tung *et al.* 2000; Sourirajan and Lichten 2008; Acosta *et al.* 2011; Argunhan *et al.* 2017). Inhibition and nuclear exclusion of Ndt80 by the checkpoint prevents the wave of meiotic induction of Clb1 required for entry into meiosis I (Wang *et al.* 2011). In addition, stabilization of Swe1 by MRC action also maintains Cdc28^CDK^ inhibited by Tyr19 phosphorylation (Leu and Roeder 1999). In sum, the lack of Clb1 induction together with the inhibitory phosphorylation of Cdc28 restrains prophase I exit by keeping in check CDK activity levels.

Most of DNA meiotic transactions occur in the context of highly specialized chromosome and chromatin structures. Chromatin dynamics can be modulated by several processes, including posttranslational modification (PTM) of histones and incorporation of histone variants. Among the myriad of histone PTMs described to date, a meiotic function has been ascribed to a number of them (Brachet *et al.* 2012; Wang *et al.* 2017). In particular, H3K79 methylation and H4K16 acetylation are involved in the budding yeast MRC (Ontoso *et al.* 2013; Cavero *et al.* 2016). Much less is known about the meiotic functional contribution of histone variants; in particular, one of the most prominent such as H2A.Z, a variant of the canonical histone H2A conserved in evolution from yeast to human.

In vegetative yeast cells, H2A.Z is involved in multiples processes, including transcription regulation (both positively and negatively), maintenance of genome stability and chromatin silencing (Billon and Cote 2013; Weber and Henikoff 2014). H2A.Z is preferentially found in the vicinity of promoters at nucleosomes flanking a nucleosome-depleted region containing the transcription start site (Raisner *et al.* 2005). Nevertheless, not all the functions of H2A.Z are necessarily related to transcription; for example, H2A.Z is also deposited at persistent DSBs promoting their anchorage to the nuclear periphery and stimulating resection (Kalocsay *et al.* 2009; Adkins *et al.* 2013; Horigome *et al.* 2014). The incorporation of H2A.Z to chromatin is carried out by the SWR1 complex, which utilizes the energy of ATP hydrolysis to exchange canonical H2A-H2B by H2A.Z-H2B dimers in particular nucleosomes (Krogan *et al.* 2003; Kobor *et al.* 2004; Mizuguchi *et al.* 2004).

The number of studies addressing the role(s) of H2A.Z during meiosis is much scarce, although H2A.Z also appears to perform meiotic functions in several model organisms. In *Arabidopsis thaliana*, H2A.Z is associated to meiotic recombination hotspots and colocalizes with chromosomal foci of the Dmc1 and Rad51 recombinases; moreover, meiocytes from the *arp6* mutant (lacking a component of the SWR1 complex) show reduced number of Dmc1, Rad51 and Mlh1 foci suggesting a role for H2A.Z in the formation and/or processing of meiotic DSBs (Choi *et al.* 2013). Meiotic gene expression is also altered in the *arp6* mutant of *A. thaliana* (Qin *et al.* 2014). During mouse spermatogenesis, H2A.Z is first detected at pachytene, but excluded from the sex-body, where it accumulates at later stages. Based on the dynamics of chromosomal distribution during mammalian spermatogenesis a role for H2A.Z in meiotic sex chromosome inactivation has been proposed (Greaves *et al.* 2006; Ontoso *et al.* 2014). Recently, a transcription-independent function of H2A.Z in meiotic DSB generation by modulating chromosomal architecture in the fission yeast *Schizosaccharomyces pombe* has been reported (Yamada *et al.* 2018).

In contrast to most organisms where the absence of H2A.Z is not compatible with life, the *htz1* deletion mutant in *S. cerevisiae* (lacking H2A.Z) is viable, allowing us to directly assess its meiotic functional impact. In most cases the role of H2A.Z in other organisms has been inferred indirectly by analyzing mutants of the SWR1 complex or by cytological observations. In this work, we demonstrate that H2A.Z is important for meiosis in the budding yeast *S. cerevisiae*. We show that the *htz1* mutant displays impaired meiotic progression and sporulation and that spore viability is compromised, although meiotic interhomolog recombination does not appear to be strongly affected. The landscape of gene expression during meiotic prophase is substantially altered in the absence of H2A.Z, likely contributing to at least some of the *htz1* meiotic phenotypes. Finally, we report that H2A.Z also functions during the meiotic checkpoint response induced by the *zip1* mutant impacting on the regulators of the prophase to meiosis I transition, such as the Ndt80 transcription factor and the CDK inhibitory kinase Swe1. Our study reveals the existence of novel functional connections between these cell cycle regulators.

## Materials and Methods

### Yeast strains

Yeast strains genotypes are listed in Table S1. All the strains, except the ones used in Figure 1F and Figure S1, are isogenic to the BR1919 background (Rockmill and Roeder 1990). The *htz1::hphMX4*, *swr1::natMX4*, *swr1::hphMX4*, *spo11::natMX4*, *sum1::natMX4*, *mer3:hphMX4*, *swe1::natMX4, [hta1-htb1]::kanMX6* and *[hta2-htb2]::natMX4* gene deletions were made using a PCR-based approach (Longtine *et al.* 1998; Goldstein and McCusker 1999). The *htz1::URA3* deletion was made using the pTK17 plasmid digested with *Hin*dIII-*Sal*I (Santisteban *et al.* 2000). The *zip1::LYS2*, *mek1::kanMX6*, *ddc2::TRP1*, *sml1::kanMX6*, *spo11::ADE2*, *swe1::LEU2* and *rad51::natMX4* gene deletions were previously described (Leu and Roeder 1999; San-Segundo and Roeder 1999; Refolio *et al.* 2011; Ontoso *et al.* 2013; Herruzo *et al.* 2016). *HTZ1-GFP* and *MIH1-GFP* were made by PCR using pKT127 (Sheff and Thorn 2004) and pFA6a-kanMX6-GFP (Longtine *et al.* 1998), respectively. The *P_GAL1_-ZIP1-GFP* and *P_GDP1_-GAL4(848).ER* constructs were obtained from Amy Macqueen (Wesleyan University, CT, USA) (Voelkel-Meiman *et al.* 2012). Strains carrying Swe1 tagged with 3 copies of the MYC epitope at the N terminus and strains carrying *ZIP1-GFP* have been previously described (Leu and Roeder 1999; White *et al.* 2004). The kinase-dead *swe1-N584A* allele was generated using the *delitto perfetto* approach (Stuckey *et al.* 2011). Strains carrying the *cdc28-AF* mutation, in which Thr18 and Tyr19 of Cdc28 have been changed to alanine and phenylalanine, respectively, were generated by transformation with the plasmid pR2042 digested with *Blp*I (Leu and Roeder 1999). The *htb1-Y40F* mutant strains in which the Y40 of histone H2B has been mutated to phenylalanine carry the deletion of the *HTA1-HTB1* and *HTA2-HTB2* genomic loci and a centromeric plasmid (pSS348) expressing *HTA1-htb1-Y40F*. These strains were generated as follows. A diploid heterozygous for *[hta1-htb1]::kanMX6* and *[hta2-htb2]::natMX4* containing the *URA3*-based pSS345 plasmid expressing wild-type *HTA1-HTB1* was transformed with the *TRP1*-based pSS347 or pSS348 plasmids expressing wild-type *HTA1-HTB1* (as control) or *HTA1-htb1-Y40F,* respectively (see plasmid construction below). These diploids were sporulated and Ura-Trp+ haploid segregants harboring *[hta1-htb1]::kanMX6* and *[hta2-htb2]::natMX4* genomic deletions and the pSS347 or pSS348 plasmid as the only source for H2A-H2B or H2A-H2BY40F, respectively, were selected. In all cases, gene deletions, mutations and tagging in haploid strains were made by direct transformation with PCR-amplified cassettes and/or digested plasmids, or by genetic crosses and sporulation (always in an isogenic background) followed by selection of the desired segregants. Diploids were made by mating the corresponding haploid parents and isolation of zygotes by micromanipulation.

**Figure 1.**
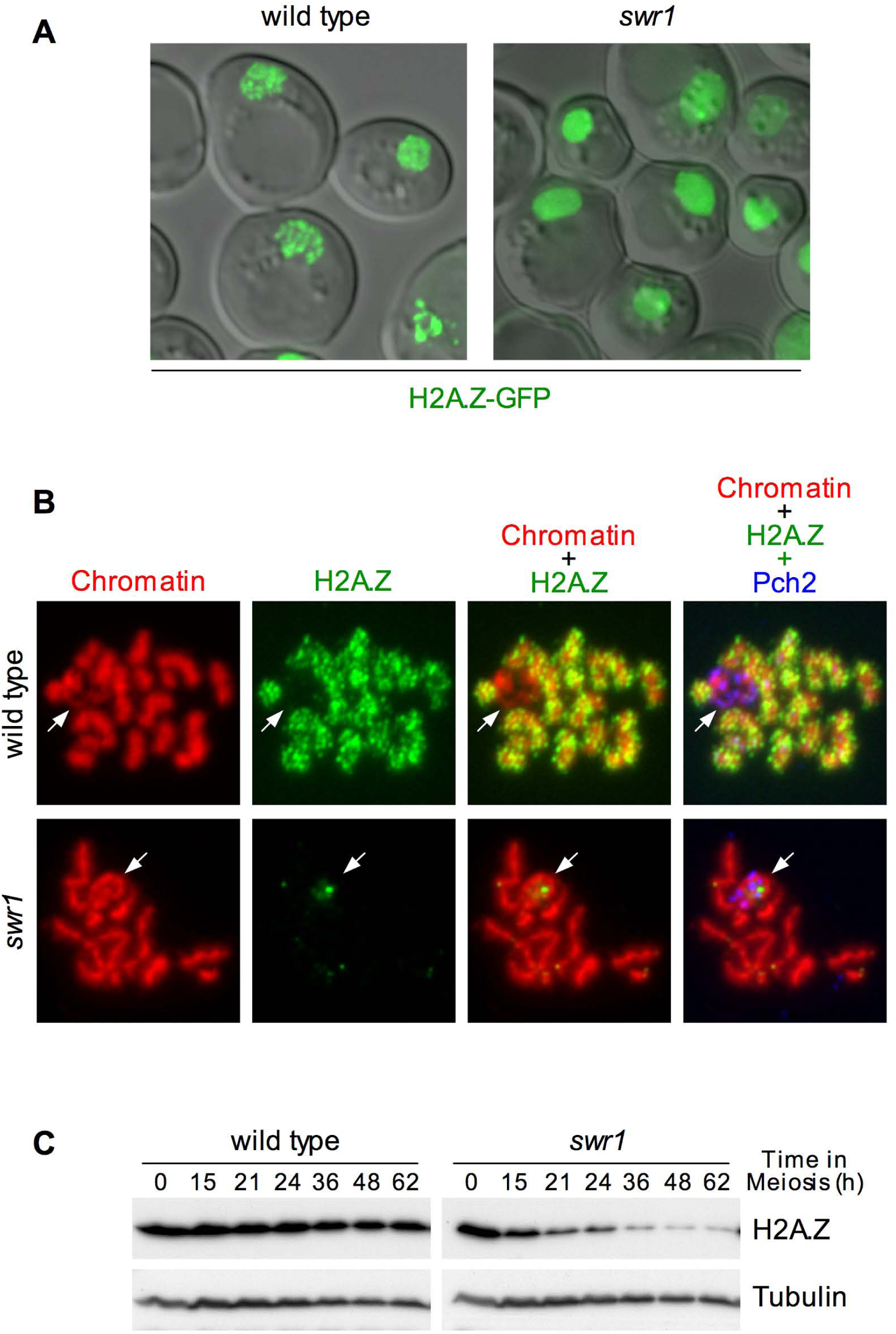
Localization of H2A.Z during meiotic prophase depends on SWR1. (A) Representative images of wild type and *swr1* live cells, at 15 hours after meiotic induction (peak of prophase I), expressing *HTZ1-GFP*. (B) Immunofluorescence of spread meiotic chromosomes from wild type and *swr1* stained with DAPI (red) to visualize chromatin, anti-GFP (green) to detect H2A.Z, and anti-Pch2 (blue) to mark the nucleolar region (arrow). (C) Western blot analysis of H2A.Z production during meiosis detected with anti-GFP antibodies. Tubulin was used as a loading control. Strains are DP840 (*HTZ1-GFP*) and DP841 *(swr1 HTZ1-GFP*).

### Plasmids

The plasmids used in this work are listed in Table S2. The 2μ-based high-copy pSS248 plasmid contains the meiosis-specific *HOP1* promoter driving the expression of GFP. In-frame cloning of a gene ORF after the GFP in pSS248 leads to overproduction of the GFP-fusion specifically during meiotic prophase. pSS248 was constructed as follows. First, the *HOP1* promoter (650 bp) was amplified from genomic DNA and cloned into the *Bgl*II-*Pac*I sites of pFA6a-kanMX6-GAL1-GFP (Longtine *et al.* 1998) replacing the *GAL1* promoter by the *HOP1* promoter to generate pSS232. Then, the *P_HOP1_-GFP* fragment from pSS232 was amplified by PCR with oligonucleotides containing the appropriate restriction sites and cloned into *Spe*I-*Not*I of pYES2 (Invitrogene) to replace *P_GAL1_* by *P_HOP1_-GFP* generating pSS248. The pSS265 plasmid to overexpress *MIH1* during meiosis was constructed by PCR amplification of the *MIH1* ORF flanked by *Not*I-*Spe*I sites and cloning into *Not*I-*Xba*I of pSS248. For meiotic overexpression of *BDF1* the ORF flanked by *Not*I-*Sph*I sites was amplified by PCR and cloned into the same sites of pSS248 to generate pSS354. Plasmid pSS263 was generated by cloning a 2.7-kb *Not*I-*Sal*I fragment from pSS200 (=p1-1) (Pak and Segall 2002) containing *NDT80* plus the promoter and 3’UTR regions into the same sites of the high-copy vector pRS426. The *HTA1-HTB1* genomic region containing the genes encoding histones H2A and H2B expressed from a common divergent promoter including also 285 and 540 bp of the flanking 3’UTR sequences was amplified by PCR from genomic DNA and cloned into the *Bam*HI-*Sal*I sites of the centromeric vectors pRS316 and pRS314 to generate plasmids pSS345 and pSS347, respectively. The *htb1-Y40F* mutation was introduced by site-directed mutagenesis of pSS347 to generate pSS348. All constructs were verified by sequencing. Oligonucleotide sequences are available upon request.

### Meiotic time courses, sporulation efficiency and spore viability

For meiotic time courses, BR strains were grown in 3.5 ml of 2xSC medium for 20 - 24 hours (2% Glucose, 0.7% Yeast Nitrogen Base without amino acids, 0.05% Adenine and Complete Supplement Mixture from Formedium at twice the particular concentration indicated by the manufacturer), then transferred to YPDA (2.5 ml) and incubated to saturation for additional 8 hours. Cells were harvested, washed with 2% potassium acetate (KAc), resuspended into 2% KAc (10 ml) and incubated at 30°C with vigorous shaking to induce meiosis and sporulation. Both YPDA and 2% KAc were supplemented with 20 mM adenine and 10 mM uracil. The culture volumes were scaled-up when needed. To score meiotic nuclear divisions, samples were taken at different time points, fixed in 70% Ethanol, washed in PBS and stained with 1 mg/ml DAPI for 15 min. At least 300 cells were counted at each time point. Meiotic time courses were repeated several times. To induce *ZIP1-GFP* from the *P_GAL1_* promoter in strains expressing *GAL4.ER*, 1 mM β-estradiol (Sigma E2257; dissolved in ethanol) was added to the cultures. Sporulation efficiency was quantitated by microscopic examination of asci formation after 3 days on sporulation plates. Both mature and immature asci were scored. At least 300 cells were counted for every strain. Spore viability was assessed by tetrad dissection. At least 144 spores were scored for every strain.

### Western blotting

Total cell extracts were prepared by TCA precipitation from 5-ml aliquots of sporulation cultures as previously described (Acosta *et al.* 2011). Analysis of Mek1 phosphorylation using Phos-tag gels was performed as reported (Ontoso *et al.* 2013). The antibodies used are listed in Table S3. The ECL or ECL2 reagents (ThermoFisher Scientific) were used for detection. The signal was captured on films and/or with a ChemiDoc XRS system (Bio-Rad).

### Fluorescence microscopy

Immunofluorescence of chromosome spreads was performed essentially as described (Rockmill 2009). For analysis of spindle formation by whole-cell immunofluorecence, the following protocol was used. Cells from meiotic cultures (1.5 ml) were fixed with 3.7% formaldehyde for 45 minutes, washed twice with solution A (1.2 M Sorbitol, 0.05 M KH_2_PO_4_) and resuspended into the same solution containing 0.1 mg/ml 20T Zymolyase, 0.1% Glusulase and 0.001% β-mercaptoethanol. Samples were incubated at 37°C for 20-30 minutes monitoring spheroplast formation. After two washes with ice-cold Solution A, cells were resuspended into 50 μl of this solution. 25 μl were deposited onto a polylysine-coated 8-well glass slide and let stand for 30 minutes. Liquid was carefully aspirated and the slide was submerged into-20°C methanol for 6 minutes and −20°C acetone for 30 seconds using a Coplin jar. The wells were successively rinsed with 1% BSA in PBS, 1% BSA 0.1% NP-40 in PBS (twice) and 1% BSA in PBS, and incubated overnight with the anti-tubulin antibody in 1% BSA-PBS at 4°C. Wells were then rinsed as described above, incubated with the secondary antibody for 2 hours at room temperature and rinsed again. A drop of Vectashield containing DAPI (Vector Laboratories, H-1200) was deposited and extended with a coverslip sealed with nail polish. The antibodies used are listed in Table S3. Images of spreads and fixed whole cells were captured with a Nikon Eclipse 90i fluorescence microscope controlled with MetaMorph software and equipped with a Hammamatsu Orca-AG CCD camera and a PlanApo VC 100×1.4 NA objective. Images of fluorescent spores as well as *ZIP1-GFP* and *HTZ1-GFP* live cells were captured with an Olympus IX71 fluorescence microscope equipped with a personal DeltaVision system, a CoolSnap HQ2 (Photometrics) camera and 100x UPLSAPO 1.4 NA objective. For Zip1-GFP and Htz1-GFP, stacks of 10 planes at 0.4 μm intervals were captured. Maximum intensity projections of deconvolved images were generated using the SoftWorRx 5.0 software (Applied Precisions). DAPI images were collected using a Leica DMRXA fluorescence microscope equipped with a Hammamatsu Orca-AG CCD camera and a 63x 1.4 NA objective.

### Recombination frequency

To measure genetic distances in a chromosome VIII interval we used a spore-autonomous fluorescence assay in SK1 strains as previously described (Thacker *et al.* 2011). Basically, diploid SK1 cells were patched on YEP-glycerol plates, streaked on YPD plates and single colonies were inoculated into 2 ml of liquid YPD incubated at 30°C for 20 h. Cells were transferred to 10 ml of YPA (1% yeast extract, 2% peptone, 2% KAc), incubated at 30°C for 14 h and sporulated in 10 ml of 2% potassium acetate containing 0.001% polypropylene glycol to prevent aggregation. Asci with fluorescent spores were imaged after 48 h in sporulation. Samples were sonicated for 15 sec before imaging. The “cell counter” plugin of ImageJ (http://imagej.nih.gov/ij/plugins/cell-counter.html) was used to manually score the tetrads of each type. Genetic distances (cM) were calculated using the Perkins equation: cM = (100 (6NPD + T))/(2(PD + NPD + T)), where PD is the number of parental ditypes, NPD is the number of nonparental ditypes and T is the number of tetratypes.

### Meiotic transcriptome analysis

Global analysis of gene expression during meiotic prophase was carried essentially as described (Morillo-Huesca *et al.* 2010). Briefly, gene expression profiles were determined using the “GeneChip™ Yeast Genome 2.0 Array” of Affymetrix at CABIMER Genomics Unit (Seville, Spain). Total RNA from meiotic prophase cells (15 hours after meiotic induction) was isolated using the RNeasy^®^ Midi kit (Qiagen) and its quality confirmed by Bioanalyzer^®^ 2100 (Agilent technology). Synthesis, labeling and hybridization of cDNA was performed with RNA from 3 independent cultures of each strain following Affymetrix protocols (http://www.affymetrix.com/analysis/index.affx). Probe signal intensities were extracted from the scanned images and analyzed with the GeneChip Operating Software (Affymetrix). The raw data (CEL files) were preprocessed and normalized using the Robust Multichip Average (RMA) method. Fold-change values (log2) and their FDR-adjusted p-values were calculated with Limma (Linear Models for Microarray Analysis using the *affylmGUI* interface. All the statistical analysis was performed using *R* language and the packages freely available from “Bioconductor Project” (http://www.bioconductor.org). Fold-change cutoffs were analyzed at 95% confidence levels (FDR-adjusted p-values<0.05). All data is MIAME compliant and the raw data have been deposited at the Miame compliant Gene Expression Omnibus (GEO) database at the National Center for Biotechnology Information (http://www.ncbi.nlm.nih.gov/geo/) and are accessible through accession number GSE110022. Gene ontology and functional clustering analyses were performed using DAVID tools (Database for Annotation, Visualization and Integrated Discovery) (Huang *et al.* 2007).

### Quantitative RNA analysis

The amount of mRNA was determined by real-time PCR (RT-PCR) amplification of cDNA generated by reverse transcription and RNaseH treatment (SuperScript II Reverse Transcriptase; Invitrogen) of the RNA samples obtained for the microarray hybridization analysis. Amplification of *ACT1* was used to normalize for differences in the amount of input RNA. Similar results were obtained after normalization with *NUP84* (data not shown). Primers were designed using the Primer Express software (Applied Biosystems) and their sequence is available upon request.

### Statistics

Unless specified, to determine the statistical significance of differences a two-tailed Student *t*-test was used. *P*-Values were calculated with the GraphPad Prism 5.0 software. The nature of errors bars in graphical representations and the number of biological replicates (n) is indicated in the corresponding figure legend. For analysis of statistical significance in Venn diagrams a hypergeometric test was applied.

### Data availability

All relevant data necessary for confirming the conclusions presented are within the article and Supplemental Information (GSA Figshare). Strains and plasmids are available upon request. Microarray raw data are deposited at GEO repository under GSE110022 accession number (https://www.ncbi.nlm.nih.gov/geo/query/acc.cgi?acc=GSE110022).

## Results

### H2A.Z localizes to meiotic prophase chromosomes in a SWR1-dependent manner

To investigate the localization of H2A.Z during meiotic prophase we generated a functional version of this histone variant tagged with the green fluorescent protein (GFP). Live wild-type cells observed by fluorescence microscopy 15 h after meiotic induction (peak of prophase in the BR strain background) displayed the H2A.Z-GFP signal along elongated structures likely corresponding with zygotene-pachytene chromosomes. In contrast, the *swr1* mutant showed diffused H2A.Z-GFP throughout the nucleus (Figure 1A). To explore H2A.Z localization in more detail we performed immunofluorescence of meiotic chromosome spreads (Figure 1B). In wild-type pachytene chromosomes, H2A.Z decorated all chromatin, except a particular region of the genome corresponding to the rDNA region, as demonstrated by the presence of the nucleolar-enriched Pch2 protein (Herruzo *et al.* 2016) (Figure 1B, arrows). In contrast, pachytene chromosomes of the *swr1* mutant were largely devoid of chromatin-associated H2A.Z (Figure 1B). These observations indicate that, like in vegetative cells (Krogan *et al.* 2003; Kobor *et al.* 2004; Mizuguchi *et al.* 2004), the SWR1 complex is also required for the deposition of H2A.Z into meiotic chromatin. Occasionally, discrete dots of H2A.Z-GFP accumulation could be observed in *swr1* nuclei. The nature and possible functional implication of this SWR1-independent localization of H2A.Z will be described elsewhere. Western blot analysis revealed that global levels of H2A.Z remained fairly constant throughout the whole meiotic program in the wild type; however, they were gradually diminishing in the *swr1* mutant (Figure 1C), suggesting that chromatin incorporation stabilizes H2A.Z during meiosis.

### Meiotic progression and sporulation are impaired in the *htz1* mutant

To determine whether H2A.Z plays a role in meiotic progression, we followed the kinetics of meiotic divisions by DAPI-staining of nuclei in wild type and *htz1* strains. Completion of meiotic divisions was less efficient in the *htz1* mutant compared to the wild type (Figure 2A). Likewise, sporulation efficiency and spore viability were also reduced in *htz1*, and asci morphology was altered (Figure 2B-2D). These observations imply that H2A.Z function is required for normal meiotic development. The *htz1* mutant showed a random pattern of spore death, with no predominance of four-, two- and zero-spore-viable tetrads (Figure 2E), suggesting that the reduced spore viability in *htz1* is not resulting, at least exclusively, from meiosis I non-disjunction events. We also examined crossover recombination in a chromosome VIII interval between *CEN8* and *THR1* using a microscopic fluorescence assay that is independent of spore viability (Thacker *et al.* 2011). Recombination frequency in this interval, measured as map distance (cM), was not altered in the *htz1* mutant compared to the wild type. As a control, a crossover-defective *mer3* mutant was also included in the assay (Figure 2F and Figure S1). To assess whether the inefficient meiotic progression of *htz1* was a consequence of the activation of the meiotic recombination checkpoint we combined the absence of H2A.Z with that of Spo11 (lacking recombination-initiating meiotic DSBs) and with that of Mek1 (lacking the main checkpoint effector kinase). The *htz1* delay in meiotic progression was maintained in the *htz1 spo11* and *htz1 mek1* double mutants (Figures 3A and 3B, respectively). Moreover, the dynamics of various indicators of checkpoint activity, such as Hop1-T318 phosphorylation (Herruzo *et al.* 2016) and Mek1 activation, as assessed both by Mek1 autophosphorylation (Ontoso *et al.* 2013) and phosphorylation of its H3-T11 target (Cavero *et al.* 2016; Kniewel *et al.* 2017), was similar in wild type and *htz1* (Figure 3C). These results indicate that the lower overall efficiency of meiotic divisions in *htz1* does not stem from activation of the meiotic recombination checkpoint, and it is consistent with the observation that meiotic recombination (CO) does not appear to be significantly affected in the absence of H2A.Z. To explore the possibility that the absence of H2A.Z affects meiotic entry rather than (or in addition to) meiotic progression we used *ZIP1-GFP* as a reporter for early meiotic gene expression and analyzed the percentage of cells showing nuclear fluorescence in the wild-type and *htz1* strains shortly after meiotic induction.

**Figure 2.**
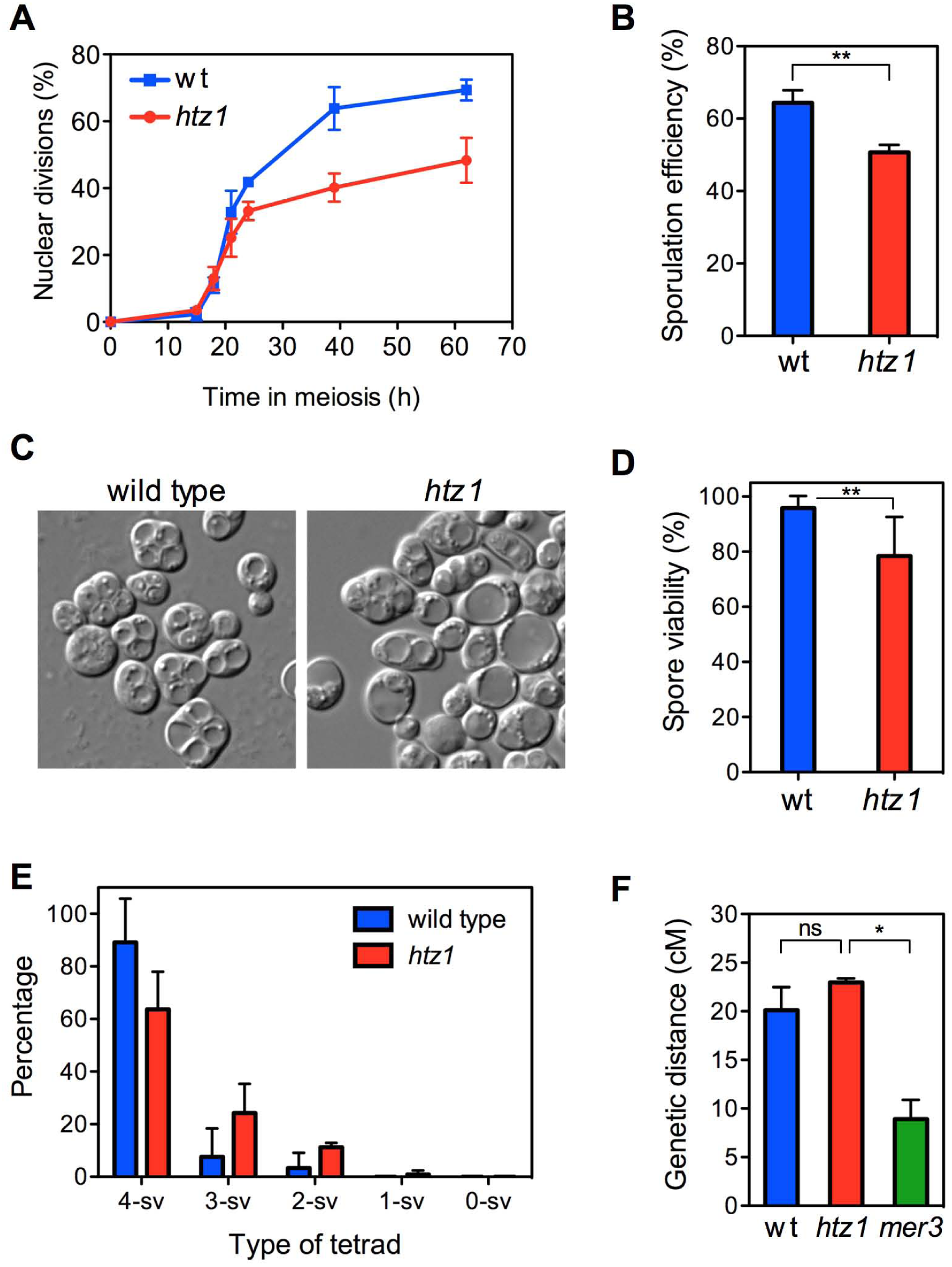
H2A.Z is required for proper meiotic development. (A) Time course analysis of meiotic nuclear divisions; the percentage of cells containing two or more nuclei is represented. Error bars: SD; n=6. (B) Sporulation efficiency, determined by microscopic counting, as the percentage of cells forming mature or immature asci after 3 days on sporulation plates. Error bars: SD; n=3. (C) Representative DIC images of asci. (D) Spore viability determined by tetrad dissection. At least 288 spores were scored for each strain. Error bars: SD; n=5. (E) Distribution of tetrad types. The percentage of tetrads with 4, 3, 2, 1 and 0 viable spores (4-sv, 3-sv, 2-sv, 1-sv and 0-sv, respectively) is represented. Error bars: SD; n=3. (F) Recombination frequency, expressed as map distance (cM), in a chromosome VIII interval (see Figure S1). Error bars: range; n=2. Strains used in (A)-(E) are DP421 (wild type) and DP630 (*htz1*). Strains used in (F) are DP969 (wild type), DP973 (*htz1*) and DP974 (*mer3*).

We found that the kinetics of appearance of Zip1-GFP fluorescence was slightly, but reproducibly, delayed in the *htz1* mutant, although eventually it reached nearly wild-type levels (Figure 3D). This observation likely reflects a delay in the onset of the meiotic program in the absence of H2A.Z and may account, at least in part, for the checkpoint-independent impaired completion of meiotic divisions in the *htz1* mutant.

**Figure 3.**
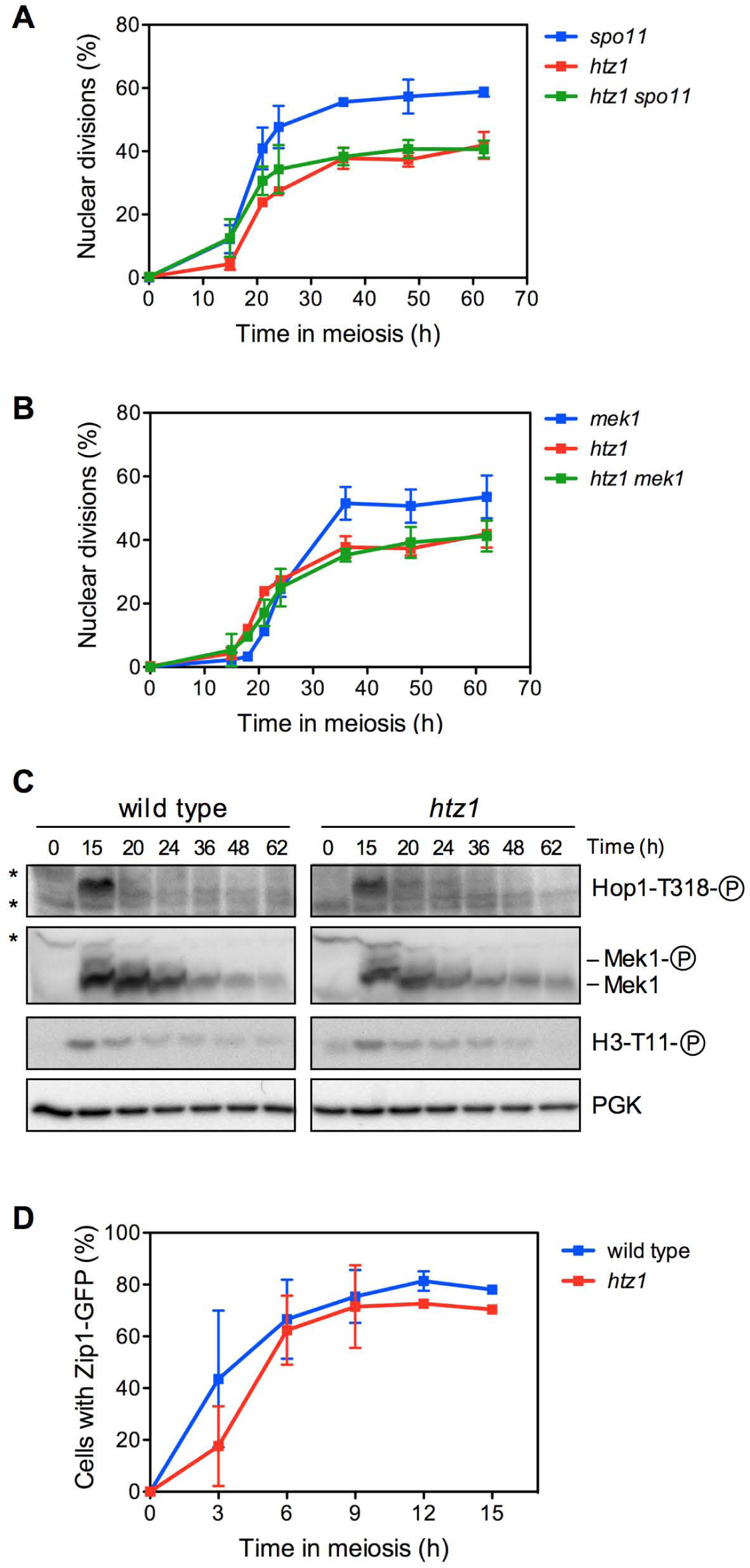
The inefficient meiotic progression of the *htz1* single mutant does not result from activation of the meiotic recombination checkpoint. (A, B) Time course analysis of meiotic nuclear divisions; the percentage of cells containing two or more nuclei is represented. Error bars: SD; n=3. (C) Western blot analysis of Hop1-T318 phosphorylation and Mek1 activity at the indicated time points in meiosis. PGK was used as a loading control. Strains in (A-C) are DP421 (wild type), DP630 (*htz1*), DP713 (*mek1*), DP1523 (*spo11*), DP1144 (*htz1 spo11*) and DP1259 (*htz1 mek1*). (D) Time course analysis of *ZIP1-GFP* induction. The percentage of cells showing Zip1-GFP nuclear fluorescence during early time points after transfer to sporulation conditions is represented. Strains are: DP437 (wild type) and DP838 (*htz1*). Error bars: SD; n=3.

### The SWR1 complex partially impairs meiosis in the absence of H2A.Z

The SWR1 complex is required to replace H2A-H2B by H2A.Z-H2B dimers at particular nucleosomes. It has been proposed that the SWR1 complex exerts a deleterious effect on chromatin integrity in the *htz1* mutant due to the attempt to replace the canonical histone H2A by H2A.Z in the absence of this histone variant creating nucleosome instability (Morillo-Huesca *et al.* 2010). As a consequence, deletion of *SWR1* (encoding the catalytic component of the SWR1 complex) totally or partially suppresses some of the multiple phenotypes of *htz1* in vegetative cells (Morillo-Huesca *et al.* 2010). Thus, we analyzed the kinetics of meiotic divisions, sporulation efficiency and spore viability in *swr1* and *htz1 swr1* mutants. We found that meiotic progression was faster (Figure S2A) and that asci formation and spore viability were somehow improved in *htz1 swr1* compared to *htz1* (Figure S2B and S2C), although they did not reach wild-type levels. The *swr1* single mutant showed an intermediate phenotype between the wild type and *htz1* mutant in meiotic progression and sporulation efficiency (Figure S2A and S2B). These observations imply that some, but not all, meiotic phenotypes of *htz1* result from the pathogenic action of SWR1 in the absence of H2A.Z. Moreover, the fact that *SWR1* deletion only partially suppresses the meiotic defects of *htz1* also supports a direct impact of H2A.Z chromatin deposition on proper meiotic development.

### Meiotic gene expression is altered in the *htz1* mutant

Several studies have demonstrated that mutation of *HTZ1* causes transcriptional misregulation during vegetative growth (Billon and Cote 2013). To assess the influence of H2A.Z on general meiotic gene expression we used whole-genome microarray analysis to compare the transcription profile of wild-type and *htz1* meiotic prophase cells (15 hours after meiotic induction). We found 611 genes showing differential expression in *htz1* compared to wild type (1.5-fold cutoff, p<0.05) (Table S4); of those, 339 genes were up-regulated and 272 down-regulated. Among the genes whose expression was increased in the absence of H2A.Z, genes encoding ribosomal proteins were on the top of the list ordered by linear-fold change (LFC) (Table S4). On the top positions of the genes whose expression was significantly down-regulated in the *htz1* mutant we found genes involved in the MEN (Mitotic Exit Network) pathway (*BFA1*, *LTE1*) and PP1 phosphatase regulators (*GIP14*, *GAC1*) (Table S4). Although there were no meiosis-specific genes among those whose mRNA levels showed a strong change, it was possible to find some genes with meiotic functions, chromatin, DNA damage response and cell-cycle related events with a LFC>1.5 (Table 1). The reduced expression of some of these genes in the *htz1* mutant was verified by RT-PCR analysis of the same mRNA samples used in the microarrays (Figure S3A). Moreover, gene ontology and clustering analyses of the genes with decreased expression showed a significant enrichment of functional categories related to both mitotic and meiotic cell cycle regulation (Table S4). On the contrary, genes with increased expression in *htz1,* cluster mainly in ribosome biogenesis, translation and metabolic processes (Table S4). Since genes encoding ribosomal proteins are rapidly repressed upon meiotic induction (Chu *et al.* 1998), this observation is consistent with the slight delay in meiosis entry of the *htz1* mutant (Figure 3D). Interestingly, 133 of 611 genes (*p*=5×10^−5^) with differential level of expression between wild type and *htz1* during meiotic prophase identified in this study overlap with those affected by *htz1* (948 genes) in mitotically growing cells (Morillo-Huesca *et al.* 2010) (Figure S3B).

Thus, these analyses revealed that the meiotic transcriptional landscape is significantly disturbed in *htz1*, suggesting that the pleiotropic phenotypes of the *htz1* mutant (aberrant morphology, inefficient meiotic development, low spore viability…) could stem from the more or less subtle alteration of multiple mechanisms.

**Table 1.** Subset of genes with decreased meiotic prophase expression in *htz1 (p*<0.05)

| GENE | LFC (>1.5) |
| --- | --- |
| Cell cycle |  |
| *BFA1* | 2.393400 |
| *LTE1* | 2.277104 |
| *GIP4* | 2.268020 |
| *BUB2* | 1.704342 |
| *MCK1* | 1.586536 |
| *MIH1* | 1.583281 |
| *CDC7* | 1.536321 |
| Meiotic genes |  |
| *RME1* | 1.852768 |
| *RPD3* | 1.799528 |
| *HFM1* | 1.683362 |
| *REC8* | 1.666158 |
| *MEK1\** | 1.599045 |
| *MRE11* | 1.597116 |
| *SKI8* | 1.570898 |
| *IME4* | 1.531359 |
| *ZIP2* | 1.511252 |
| *SPO22* | 1.504426 |
| *HOP1* | 1.499766 |
| DNA damage response |  |
| *SRS2* | 1.898810 |
| *RAD17* | 1.705357 |
| *TOF1* | 1.612276 |
| *MEC1* | 1.509767 |
| *IRC6* | 1.505446 |
| Chromatin |  |
| *SWC3* | 2.864716 |
| *SWI3* | 2.070301 |
| *RSC8* | 1.850621 |
| *SPT20* | 1.631780 |
| *HFI1* | 1.629896 |
| *CHD1* | 1.514238 |

LFC: linear fold change (>1.5)
\* $p = 0.07$

### The *zip1 htz1* mutant displays a tight checkpoint-dependent meiotic arrest

Next, we sought to explore the possible role of H2A.Z during challenged meiosis; that is, under conditions in which meiotic defects trigger the meiotic recombination checkpoint. We used the *zip1* mutant, which is defective in CO recombination and SC formation, to induce the checkpoint. The *zip1* mutant arrests in prophase I for a long period, but eventually, at late time points, a fraction of the culture completes the meiotic divisions to generate largely inviable spores (Figure 4A) (Ontoso *et al.* 2013). Strikingly, we found that meiotic progression was completely blocked in the *zip1 htz1* double mutant as most cells remained uninucleated during the whole time course (Figure 4A). This observation suggests that H2A.Z may have a role during prophase I exit because its absence, combined with that of Zip1, provokes a strong meiotic arrest.

**Figure 4.**
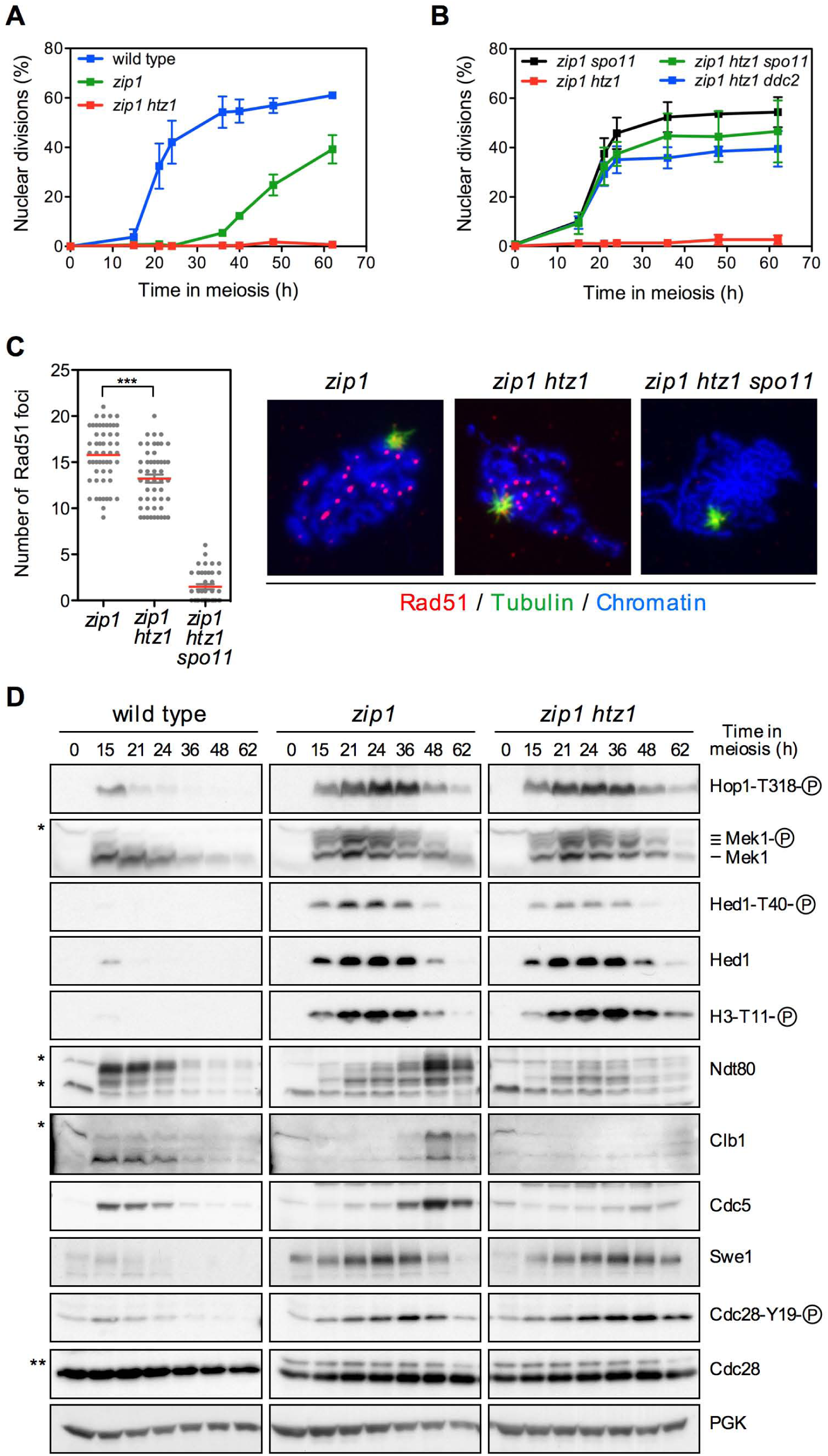
Robust checkpoint-dependent meiotic arrest *in zip1 htz1*. (A), (B) Time course analysis of meiotic nuclear divisions; the percentage of cells containing two or more nuclei is represented. Error bars: SD; n=3. Strains are DP421 (wild type), DP422 (*zip1*), DP776 (*zip1 htz1*), DP1524 (*zip1 spo11*), DP815 (*zip1 htz1 spo11*) and DP816 (*zip1 htz1 ddc2*). (C) Localization and quantification of Rad51 foci as markers for unrepaired DSBs on spread meiotic nuclei of *zip1* (DP449), *zip1 htz1* (DP776) *and zip1 htz1 spo11* (DP815) after 16 h of meiotic induction. Only prophase I nuclei, as assessed by tubulin staining, were scored. Representative images are shown. (D) Western blot analysis of the indicated molecular markers of checkpoint activity at different levels in the pathway. PGK was used as a loading control. Strains are DP421 (wild type), DP422 (*zip1*) and DP631 (*zip1 htz1*). For detection of Myc-tagged Swe1, the strains used are DP1353 (wild type), DP1354 (*zip1*) and DP1414 (*zip1 htz1*).

Like in the wild-type (Figure 1), chromatin incorporation and stability of H2A.Z also depended on SWR1 in the *zip1* mutant (Figure S4A-S4B). To determine whether the impact of *htz1* on the inability to resume meiotic progression in *zip1* was a consequence of the deleterious effect of SWR1 as explained above, we analyzed the kinetics of meiotic divisions in *zip1 swr1* and *zip1 htz1 swr1* mutants. Interestingly, like *zip1 htz1*, the *zip1 swr1* and *zip1 htz1 swr1* mutants also showed a tight meiotic block (Figure S4C). Since the *swr1* single mutant is able to complete meiosis, albeit with a small delay compared to the wild type, these results indicate that the strong meiotic arrest of *zip1 htz1*, *zip1 swr1* and *zip1 htz1 swr1* stems from the lack of H2A.Z chromatin deposition and does not result from the indirect toxic effect of SWR1 in the absence of H2A.Z.

To ascertain whether the *zip1 htz1* block was caused by the meiotic recombination checkpoint we generated the *zip1 htz1 spo11* mutant, in which meiotic DSBs are not formed (Keeney *et al.* 1997), and the *zip1 htz1 ddc2* mutant, in which meiotic recombination intermediates are not sensed (Refolio *et al.* 2011). We found that meiotic divisions and sporulation were largely restored in the *zip1 htz1 spo11* and *zip1 htz1 ddc2* mutants (Figure 4B) generating mostly dead spores (5.6% and 1.5% spore viability for *zip1 htz1 spo11* and *zip1 htz1 ddc2*, respectively; n=72), thus confirming that the meiotic prophase block in *zip1 htz1* is imposed by the meiotic recombination checkpoint.

### The *zip1 htz1* mutant does not accumulate additional unrepaired DSBs

One possible explanation for the more robust meiotic arrest of *zip1 htz1* compared to that of *zip1* is that the absence of H2A.Z may provoke additional defects that, combined to those resulting from the lack of Zip1, could lead to further hyperactivation of the meiotic recombination checkpoint and, therefore, a tighter prophase I block. To test this possibility, we used immunofluorescence of spread nuclei to analyze the presence of Rad51 foci as an indirect marker for unrepaired DSBs (Joshi *et al.* 2015) in *zip1* and *zip1 htz1* mutants. The *zip1 htz1 spo11* mutant was also included as a control for the absence of meiotic DSBs. Due to the different kinetics of meiotic progression of the strains analyzed (Figure 4A-4B), only prophase I nuclei, as assessed by the bushy morphology of tubulin staining, were scored (Figure 4C). We found that the *zip1 htz1* double mutant did not display more Rad51 foci than *zip1* (Figure 4C), suggesting that the absence of H2A.Z together with that of Zip1 does not generate more unrepaired meiotic DSBs. We also performed immunofluorescence of spread nuclei using an antibody that recognizes phosphorylated S/T-Q motifs as an additional assay for Mec1/Tel1-dependent DNA damage signaling during meiotic prophase. We found that phospho-S/T-Q foci were significantly increased in a *dmc1* mutant, used as control, that accumulates hyper-resected DSBs (Bishop *et al.* 1992), but similarly decorated prophase chromosomes of *zip1* and *zip1 htz1* (Figure S5). These observations do not favor the possibility that the accumulation of additional DNA damage is responsible for the exacerbated meiotic arrest of *zip1 htz1*.

### Dynamics of upstream checkpoint activation-deactivation is normal in *zip1 htz1*

To pinpoint what event in the *zip1*-induced MRC pathway is impacted by H2A.Z we used a battery of molecular markers to analyze checkpoint status during meiotic time courses of wild type, *zip1* and *zip1 htz1* strains (Figure 4D). Activation of the Mec1-Ddc2 sensor complex by unrepaired DSBs (and perhaps other types of meiotic defects) is one of the first events in the meiotic checkpoint pathway (Refolio *et al.* 2011; Subramanian and Hochwagen 2014). Active Mec1 phosphorylates Hop1 at various sites, including T318 (Carballo *et al.* 2008; Penedos *et al.* 2015). In the *zip1*-induced checkpoint, Hop1-T318 phosphorylation is critical to sustain activation of the Mek1 effector kinase (Herruzo *et al.* 2016), and serves as an excellent readout for Mec1 activity. Since unrepaired DSBs promote Mec1 activation, Hop1 phosphorylation has been also used as an indirect assay for DSB formation (Chen *et al.* 2015). In the wild type, there was a weak and transient phosphorylation of Hop1-T318 coincident with the peak of prophase I and ongoing recombination. In contrast, Hop1-T318 phosphorylation was very robust and sustained in the *zip1* mutant (Figure 4D), although at late time points phospho-Hop1-T318 declined coincident with completion of meiotic divisions in a fraction of the culture (Figure 4A). Remarkably, despite the tight meiotic arrest (Figure 4A), the kinetics of Hop1-T318 phosphorylation in the *zip1 htz1* double mutant was similar to that of *zip1* (Figure 4D), further supporting that the turnover of meiotic DSBs is not significantly affected by *htz1*.

We also monitored the activity of the downstream Mek1 effector kinase using three different readouts: Mek1 autophosphorylation (Ontoso *et al.* 2013), Hed1 phosphorylation at T40 (Callender *et al.* 2016) and histone H3 phosphorylation at T11 (Cavero *et al.* 2016). As shown in Figure 4D, the dynamics of Mek1 activation paralleled that of Hop1-T318 phosphorylation (that is, Mec1 activity) and, again, was similar in both *zip1* and *zip1 htz1*, except for a slight persistence of phospho-H3-T11 in *zip1 htz1* at the latest time point.

These results, together with the analysis of Rad51 foci, indicate that the robust meiotic block in *zip1 htz1* does not arise from the persistence of unrepaired recombination intermediates sustaining permanent upstream checkpoint activation.

### H2A.Z is required for reactivation of the cell cycle checkpoint targets

We next analyzed the downstream targets that are inhibited by the checkpoint to prevent cell cycle progression while recombination and/or synapsis defects persist. In particular, we examined the production of various meiosis I-promoting factors: the Ndt80 transcriptional inductor, the Clb1 cyclin and the Cdc5 polo-like kinase (Acosta *et al.* 2011). In addition, we also monitored the levels of the Swe1 kinase and its activity: the inhibitory phosphorylation of Cdc28 (CDK) at tyrosine 19 (Leu and Roeder 1999). In the wild type, after the recombination process is completed and the transient activation of Mek1 disappears, the program for meiosis I entry is turned on with the production of Ndt80, Clb1 and Cdc5, as well as the reduction of the inhibitory phosphorylation at Y19 of Cdc28 (Figure 4D). In the *zip1* mutant, the induction of Ndt80, Clb1 and Cdc5 were significantly delayed and high levels of the Swe1 kinase promoting Cdc28-Y19 phosphorylation persisted for as long as Mek1 was active. However, as Mek1 activation eventually declined, Ndt80 and Cdc5 were induced, and Swe1 and phospho-Cdc28-Y19 diminished, thus sustaining entry into meiosis I of at least a fraction of the cells (Figure 4A, 4D). In contrast, we found that although Mek1 was down-regulated in *zip1 htz1* with similar kinetics to that in *zip1*, Ndt80, Clb1 and Cdc5 production remained largely inhibited, and Swe1 and phospho-Cdc28-Y19 levels stayed high at late time points (Figure 4D), consistent with the inability of *zip1 htz1* cells to exit prophase I (Figure 4A). These results indicate that the main cell cycle targets of the checkpoint are misregulated in the absence of H2A.Z and suggest that this impairment is responsible for the strong block in meiotic progression of *zip1 htz1*.

### HA2.Z contribution to checkpoint recovery

To determine whether H2A.Z is required to re-start meiotic cell cycle progression when the *zip1* defects that initially triggered the checkpoint are corrected we used a conditional system in which *ZIP1-GFP* expression is controlled by β-estradiol. *ZIP1-GFP* was placed under control of the *GAL1* promoter in strains producing a version of the Gal4 transcriptional regulator fused the β-estradiol receptor (Gal4[848].ER) (Benjamin *et al.* 2003; Voelkel-Meiman *et al.* 2012). As depicted in Figure 5A, meiotic cultures of both wild-type and *htz1* strains were initiated without β-estradiol; that is, in the absence of Zip1, to induce the checkpoint response. After 24 h, when the cells are blocked in prophase by the checkpoint, β-estradiol was added to half of the culture and the other half was maintained in the absence of the hormone as control. Recovery from the arrest after *ZIP1* induction was monitored at the cytological level (Zip1-GFP chromosome incorporation and DAPI staining of nuclei) and at the molecular level (western blot analysis of various checkpoint markers) (Figure 5B-5D).

**Figure 5.**
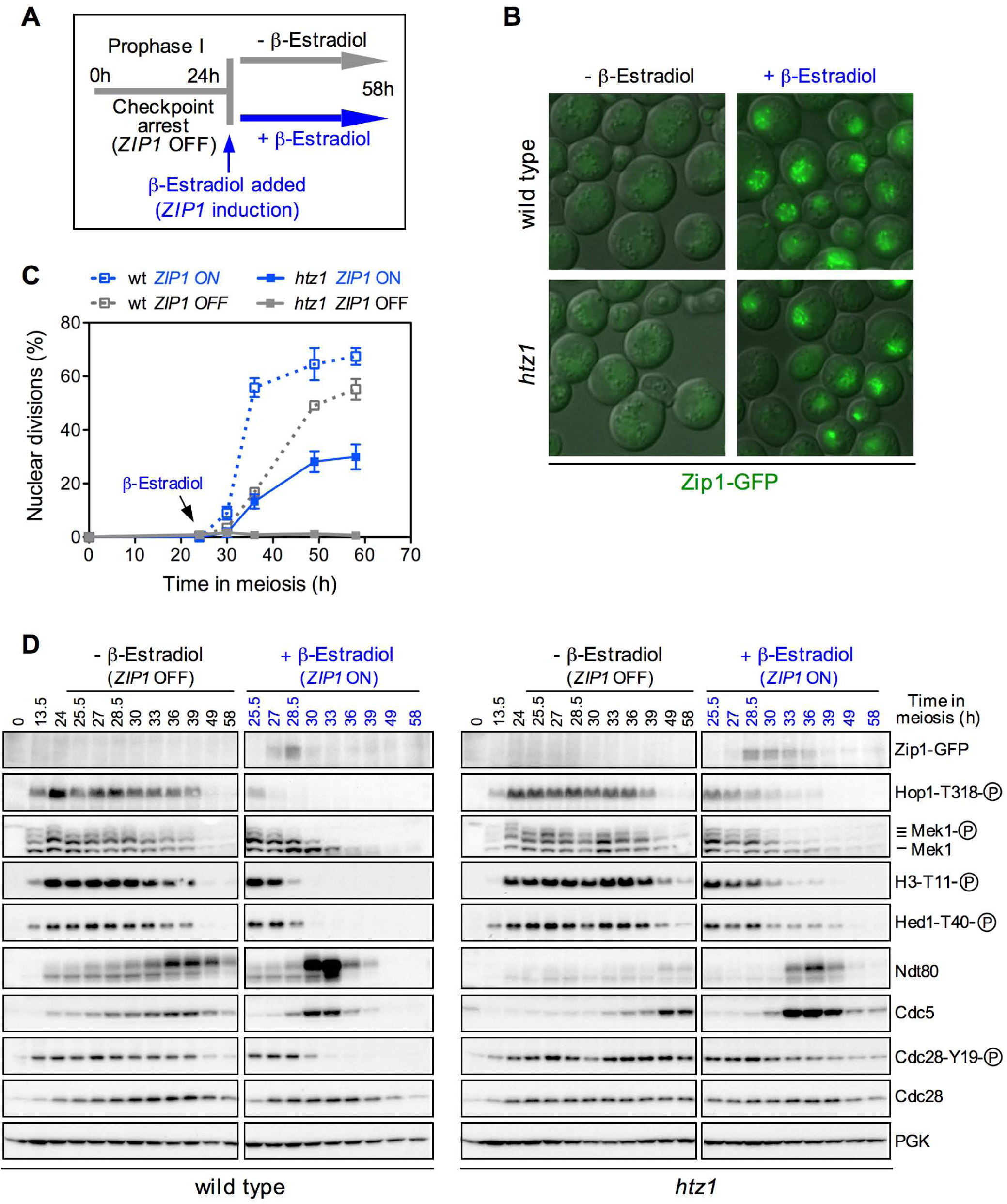
Analysis of meiotic checkpoint recovery. (A) Schematic representation of the experimental setup for conditional *ZIP1* induction in wild type (DP1185) and *htz1* (DP1186) cells containing the *GAL4-ER* transcriptional activator regulated by β-estradiol and *P_GAL1_-ZIP1-GFP*. (B) Representative fluorescence microscopy images showing SC incorporation of Zip1-GFP. Cells were imaged 3 hours after β-estradiol addition. (C) Time course analysis of meiotic nuclear divisions; the percentage of cells containing two or more nuclei is represented. The arrow indicates β-estradiol addition (blue lines and symbols). Error bars: range; n=2. (D) Western blot analysis of the indicated molecular markers of checkpoint activity. PGK was used as a loading control.

In the absence of β-estradiol (“*ZIP1* OFF”), the checkpoint was activated in the wild type as shown by the prominent H3-T11 and Hed1-T40 phosphorylation, but eventually the phosphorylation of these markers decreased concomitant with Ndt80 activation, Cdc5 production and Cdc28-Y19 dephosphorylation (Figure 5D); thus sustaining meiotic progression (Figure 5C). Note that, for unknown reasons, the meiotic delay induced by the checkpoint in this “*ZIP1* OFF” situation is less pronounced than in a *zip1* mutant (Figure 4A), perhaps due to a leaky, but undetectable, expression of *GAL1-ZIP1* even in the absence of β-estradiol. In the *htz1* mutant without β-estradiol, the checkpoint was also heavily activated but, albeit with a slightly slower kinetics, the levels of H3-T11 and Hed1-T40 phosphorylation were also finally reduced. However, like in *zip1 htz1* mutants, Ndt80 production was not induced and Cdc28-Y19 remained phosphorylated at late points (Figure 5D); as a consequence, meiotic progression was robustly blocked (Figure 5C). Thus, this “*ZIP1* OFF” situation phenocopies *ZIP1* deletion in *htz1* (Figure 4A-4D).

When β-estradiol was added, *ZIP1-GFP* expression was induced, and 3 h after hormone addition Zip1-containing chromosomes were detected in nuclei of both wild type and *htz1* (Figure 5B). *ZIP1-GFP* induction was slightly less efficient in the *htz1* mutant (Figure 5D), perhaps due to the effect of H2A.Z on *GAL1* promoter regulation (Santisteban *et al.* 2000). In the wild type, the checkpoint was rapidly turned off upon Zip1 production: Mek1 signaling drastically disappeared, Ndt80 and Cdc5 were sharply induced and Cdc28-Y19 phoshosphorylation was erased (Figure 5D; “*ZIP1* ON”). Consistently, prophase-arrested wild-type cells immediately underwent meiotic divisions after *ZIP1* expression (Figure 5C; “*ZIP1* ON”). In the *htz1* mutant the checkpoint was also down-regulated upon *ZIP1* induction, but with a slower kinetics than that of the wild type. Consistently, a fraction of *htz1* cells resumed meiotic divisions (Figure 5C, “*ZIP1* ON”); thus, H2A.Z is not essential to re-start meiotic cell cycle progression when the defects that triggered the checkpoint are resolved, but contributes to an efficient recovery from the cell-cycle arrest.

### *NDT80* overexpression partially alleviates *zip1 htz1* meiotic arrest

Since *zip1 htz1* shows a dramatic reduction in Ndt80 levels, and Cdc5 production is also impaired (Figure 4D), we examined whether an artificial increase in *CDC5* and *NDT80* expression could restore meiotic progression in *zip1 htz1*. As reported (Acosta *et al.* 2011), *CDC5* overexpression from a high-copy plasmid partially suppressed the meiotic delay of the *zip1* single mutant (Figure 6A); however, it had little effect on *zip1 htz1* (Figure 6B). In contrast, *NDT80* overexpression did promote more efficient meiotic progression in both *zip1* and *zip1 htz1* (Figure 6A-6B). These observations indicate that, in part, the strong meiotic block of *the zip1 htz1* mutant results from the drastic reduction in Ndt80 production and suggest that the relevant Ndt80-dependent event responsible for the arrest is not the inability to efficiently activate *CDC5* expression.

**Figure 6.**
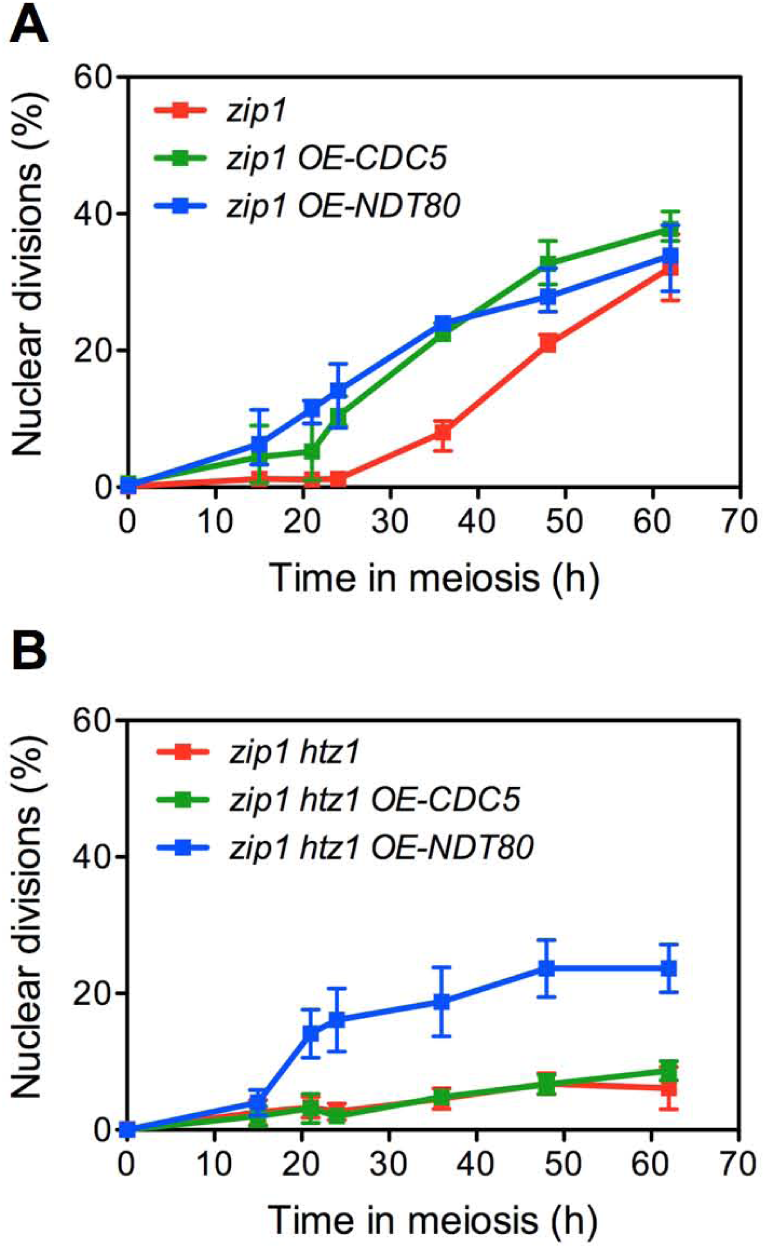
*NDT80* overexpression partially suppresses *zip1 htz1* meiotic arrest. (A), (B) Time course analysis of meiotic nuclear divisions; the percentage of cells containing two or more nuclei is represented. Strains are DP422 (*zip1*) in (A) and DP1017 (*zip1 htz1*) in (B), transformed with vector alone (pRS426) or with high-copy plasmids overexpressing *CDC5* (pJC29) or *NDT80* (pSS263), denoted as *OE-CDC5* and *OE-NDT80*, respectively. Error bars: SD; n=3.

### Deletion of *SWE1*, but not mutation of Cdc28-Y19, suppresses the *zip1 htz1* meiotic block

We have found that the levels of both the Swe1 kinase and the phosphorylation of its target, Cdc28-Y19, remain high at late time points in the *zip1 htz1* meiotic cultures. To assess the relevance of Cdc28-Y19 inhibitory phosphorylation to impose the tight *zip1 htz1* meiotic arrest (Figure 7A), we generated three situations in which this phosphorylation event is either abolished or drastically reduced (Figure 7B): 1) *SWE1* deletion, 2) *cdc28-AF* mutation (carrying the threonine 18 and tyrosine 19 of Cdc28 changed to alanine and phenylalanine, respectively), and 3) overexpression of the *MIH1* gene from the prophase I-specific *HOP1* promoter in a high-copy plasmid (Figure S6A).

Remarkably, deletion of *SWE1* conferred a notable suppression of the *zip1 htz1* meiotic arrest (Figure 7C) although it did not reach wild-type kinetics; however, the elimination of Cdc28-Y19 phosphorylation by other means, such as Mih1 overproduction or *cdc28-AF* mutation had none or only a subtle effect on meiotic progression as most cells remained uninucleated (Figure 7C); only about 10% of *zip1 htz1 cdc28-AF* cells segregated their nuclei. In contrast, *MIH1* overexpression or *cdc28-AF* mutation did accelerate meiotic progression in a *zip1* single mutant (Figure S6B). A kinase-dead *swe1-N584A* allele (Harvey *et al.* 2005) conferred the same suppression of the checkpoint meiotic arrest as the *SWE1* deletion both in *zip1* and *zip1 htz1* strains (Figure S6C), ruling out the possibility of a direct inhibitory effect exerted by the physical interaction of Swe1 with CDK independent of Tyr19 phosphorylation. Thus, these results strongly suggest that the Swe1 kinase must impact an additional mechanism, independent of CDK phosphorylation, which is particularly relevant in the absence of H2A.Z to maintain the *zip1*-induced checkpoint arrest.

**Figure 7.**
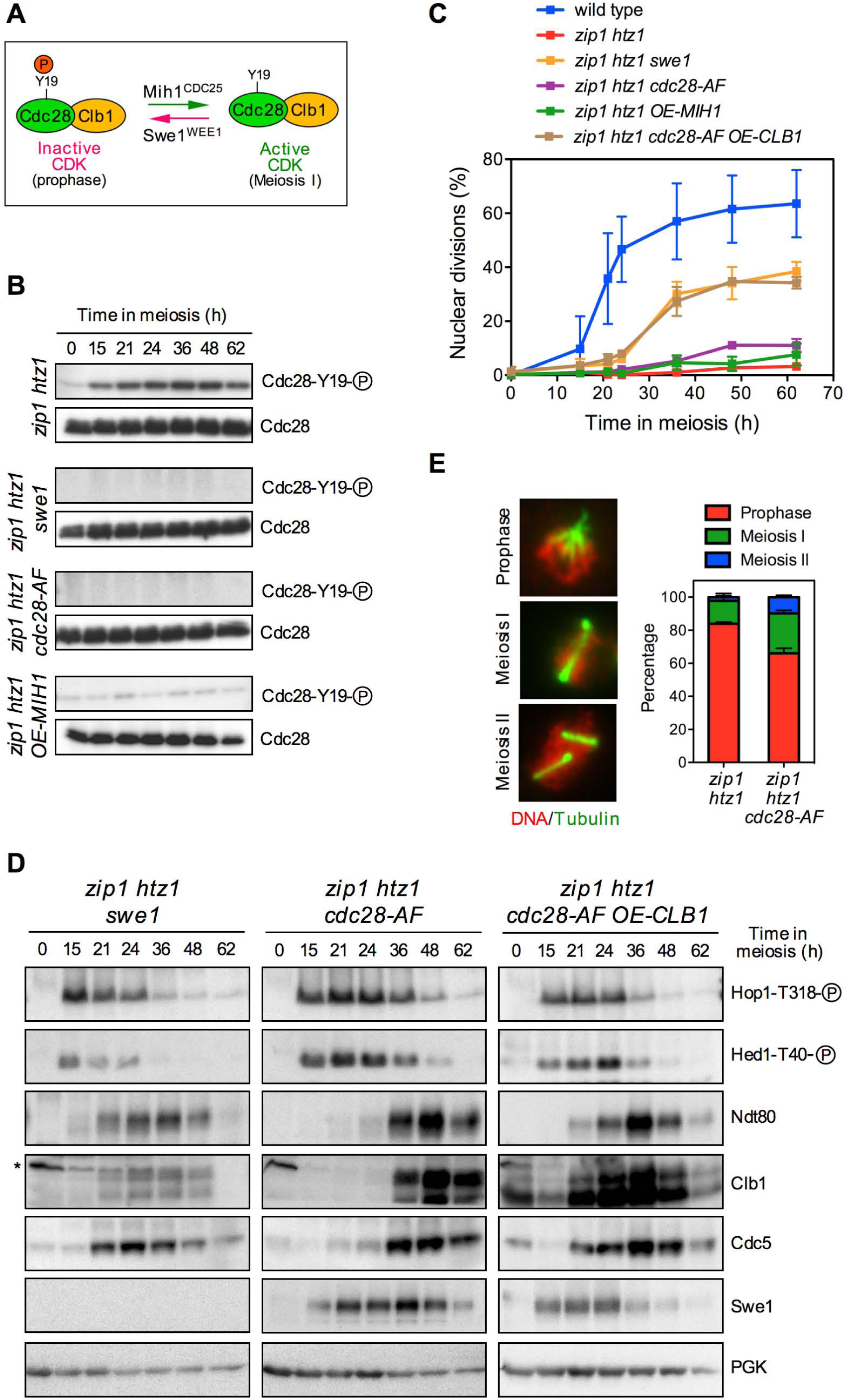
Impact of Cdc28-Y19 phosphorylation and Clb1 levels on *zip1 htz1* meiotic arrest. (A) Schematic representation of the regulation of CDK activity by Cdc28-Y19 phosphorylation controlled by the opposite action of the Swe1 kinase and the Mih1 phosphatase. (B) Western blot analysis of Cdc28-Y19 phosphorylation in the indicated strains. Total Cdc28 is also shown as control. (C) Time course analysis of meiotic nuclear divisions; the percentage of cells containing two or more nuclei is represented. Error bars: SD; n=3. (D) Western blot analysis of the indicated molecular markers of checkpoint activity. Swe1 was detected with anti-myc antibodies. PGK was used as a loading control. Strains in (B-D) are: DP1353 (wild type), DP1414 (*zip1 htz1*), DP1113 (*zip1 htz1 swe1*) and DP1416 (*zip1 htz1 cdc28-AF*). To overexpress *MIH1* and *CLB1*, the DP1414 and DP1416 strains were transformed with high-copy plasmids pSS265 (*OE-MIH1*) and pR2045 (*OE-CLB1*), respectively. (E) Whole-cell immunofluorescence using anti-tubulin antibodies in *zip1 htz1* (DP1017) and *zip1 htz1 cdc28-AF* (DP1154) cells at 48 hours in meiosis. Representative nuclei of prophase, meiosis I and meiosis II stages are shown. The quantification is presented in the graph. 169 and 119 nuclei were scored for *zip1 htz1* and *zip1 htz1 cdc28-AF*, respectively. Error bars: range; n=2.

#### *CLB1* overexpression restores meiotic progression in *zip1 htz1 cdc28-AF*

To further explore the checkpoint response in *zip1 htz1* and the effect of CDK phosphorylation we analyzed by western blot various molecular markers in the *swe1* and *cdc28-AF* mutants. According with its meiotic progression (Figure 7C), the checkpoint was deactivated in *zip1 htz1 swe1*, as manifested by the disappearance of phospho-Hop1-T318 and phospho-Hed1-T40. Concurrently, the meiosis I-promoting factors Ndt80, Clb1 and Cdc5 were produced, albeit with slower kinetics than in the wild type (Figure 7D).

Like in *zip1 htz1*, upstream checkpoint signals were also down-regulated in *zip1 htz1 cdc28-AF*; in contrast, Ndt80, Clb1 and Cdc5 accumulated at higher levels at later time points in this mutant (Figure 7D and Figure S6D). The presence of meiosis I-promoting factors suggests that the *zip1 htz1 cdc28-AF* triple mutant is proficient to undergo the prophase to meiosis I transition, but does not efficiently complete chromosome segregation. Indeed, about 40% of *zip1 htz1 cdc28-AF* cells assembled meiotic spindles at late time points (Figure 7E) despite their marked impairment to undergo meiotic divisions (Figure 7C).

Notably, *CLB1* overexpression from a high-copy plasmid restored substantial meiotic progression in *zip1 htz1 cdc28-AF* phenocopying *zip1 htz1 swe1* (Figure 7C-7D). In sum, these observations suggest that, in addition to phosphorylate Cdc28 at tyrosine 19 to prevent exit from prophase I, Swe1 regulates timing and/or abundance of Clb1 production to restrain meiotic progression in *zip1 htz1* at a later stage in meiotic development.

## Discussion

The H2A.Z histone variant is a ubiquitous determinant of chromatin structure playing crucial roles in genome stability and gene expression in mitotically dividing eukaryotic cells. However, only a limited number of studies in a few model organisms have addressed the relevance of H2A.Z in meiosis, often using indirect approaches. In this article, we have focused on the direct functional contribution of H2A.Z during meiosis in the budding yeast *S. cerevisiae*, a widely used model system for meiotic studies.

### H2A.Z is required for proper meiotic development

We report here that the *htz1* mutant of *S. cerevisiae* lacking the H2A.Z histone completes the meiotic program albeit less efficiently than the wild type. The *htz1* mutant shows delayed entry into meiosis, impaired sporulation and reduced spore viability indicating that H2A.Z is required to sustain accurate meiosis. The persistence of recombination intermediates or incomplete synapsis triggers the so-called pachytene checkpoint or meiotic recombination checkpoint (MRC) that delays meiotic progression. We found that checkpoint elimination by deleting *MEK1* or abolishing DSB formation by deleting *SPO11* do not restore normal levels of meiotic nuclear divisions in *htz1* indicating that the faulty events resulting in impaired completion of meiotic development are not sensed by the MRC and likely do not involve recombination.

In fission yeast, H2A.Z participates in the initiation of meiotic recombination by promoting the association of Spo11 and accessory proteins to chromatin (Yamada *et al.* 2018). We have found a modest reduction in the number of Rad51 foci in *zip1 htz1* compared to *zip1* (Figure 4C) that could be compatible with reduced number of initiating DSBs, although a slightly defective loading of Rad51 to DSBs in the absence of H2A.Z or a delayed onset of DSB formation cannot be ruled out. A possible role for H2A.Z in DSB generation could be also inferred from the presence of H2A.Z at promoters (at least in vegetative cells) (Raisner *et al.* 2005) where most DSBs occur in *S. cerevisiae* (Pan *et al.* 2011). However, our results suggest that, in budding yeast, the functional contribution of H2A.Z to DSB formation, if any, is only minor: 1) dynamics of Hop1 phosphorylation at T318, which serves as an indirect reporter for meiotic DSBs, is similar in wild type and *htz1*. 2) A reduction in DSB formation provoked by the absence of H2A.Z would result in a less stringent checkpoint response; however the *zip1 htz1* double mutant displays a more robust checkpoint arrest compared to *zip1*. 3) Crossover recombination in a particular interval of chromosome VIII is not significantly affected by *htz1*. It is formally possible that recombination could be altered in other chromosomal regions and/or that CO homeostasis could compensate for a reduced number of initiating events (Martini *et al.* 2006), but this would imply at best a subsidiary function for H2A.Z in DSB formation. In sum, we do not favor the scenario in which the meiotic phenotypes of the *htz1* mutant could be solely explained by impaired DSB formation. Our genome-wide study of meiotic gene expression in the *htz1* mutant reveals that many down-regulated genes cluster in several functional categories related to mitotic and meiotic cell cycle and chromosome segregation events (Table 1 and Table S4). We propose that, in unperturbed conditions, H2A.Z is not essential to perform any critical meiotic event, but the massive transcription misregulation that occurs in the absence of this histone variant may impact on various processes resulting in a less accurate and efficient completion of the meiotic program.

### H2A.Z is essential to resume meiotic progression in the absence of Zip1

Certain chromatin modifications are crucial for checkpoint activity. Dot1-mediated trimethylation of H3K79 controls Pch2 chromosomal distribution and sustains Hop1 phosphorylation and the ensuing Mek1 activation in *zip1* mutants. As a consequence, deletion of *DOT1* or mutation of H3K79 suppresses the meiotic arrest/delay of *zip1* (San-Segundo and Roeder 2000; Ontoso *et al.* 2013). The Sir2 histone deacetylase is also essential for the *zip1*-induced MRC. One of the main targets of Sir2 is acetylated H4K16. In *zip1 sir2* mutants, as well as in *zip1 H4-K16Q* mutants (mimicking constitutive H4K16 acetylation), the *zip1* block is bypassed (San-Segundo and Roeder 1999; Cavero *et al.* 2016). At least in vegetative cells, Dot1 and the SIR complex collaborate with H2A.Z in delimiting the boundaries between euchromatin and telomeric heterochromatin (Dhillon and Kamakaka 2000; Meneghini *et al.* 2003). However, these chromatin modifications perform opposite functions in the MRC; while in *zip1 dot1* and *zip1 sir2* the meiotic delay is suppressed, *zip1 htz1* shows a stronger meiotic arrest. Our results imply that, in contrast to Dot1 and Sir2, H2A.Z is not required for checkpoint activation, but it is involved in regulation meiotic progression at least in a *zip1* mutant.

We show that the *zip1* mutant exhibits a pronounced meiotic delay, but eventually, checkpoint signaling declines, as manifested by the drop in Hop1 phosphorylation and in Mek1 activation at late time points, and at least a fraction of the culture resumes meiotic progression and completes sporulation. In principle, checkpoint deactivation and resumption of cell cycle progression can occur by two related but conceptually different phenomena: ‘checkpoint adaptation’ and ‘checkpoint recovery’. Adaptation takes place when, despite the persistence of the defects that initially triggered the checkpoint, its activity declines after a prolonged period and the cell cycle resumes without previous elimination of the damage. This process of adaptation has been extensively documented in vegetative budding yeast responding to the presence of an irreparable DSB (Pellicioli *et al.* 2001). In contrast, checkpoint recovery involves the disappearance or repair of the initial problems that stimulated the checkpoint, resulting in decreased signaling and cell cycle progression.

Previous studies suggest that the eventual checkpoint deactivation and recovery of meiotic progression in *zip1* is consequence of the disappearance of the initial defects (likely unrepaired DSBs) presumably by using the sister chromatid instead of the homolog as template for DNA repair. This is based on the observation that deletion of *RAD51*, which fundamentally compromises sister chromatid recombination (Liu *et al.* 2014; Callender *et al.* 2016), leads to a permanent arrest in *zip1* (Herruzo *et al.* 2016) (Figure S7A). In this work we report that, like *zip1 rad51*, the *zip1 htz1* double mutant also shows a tight meiotic block; however, the analysis of various checkpoint markers reveals that the cause of the arrest is different in *zip1 rad51* and *zip1 htz1*. In the *zip1 rad51* mutant, high levels of Hop1-T318 phosphorylation and Mek1 activity persist until late time points, consistent with the accumulation of unrepaired recombination intermediates that signal to the checkpoint. Consequently, Cdc28-Ty19 phosphorylation remains high and Cdc5 production is inhibited, thus explaining the meiotic arrest (Herruzo *et al.* 2016) (Figure S7B). In contrast, we show here that in *zip1 htz1*, Hop1 and Mek1 activation eventually decline with similar kinetics to that observed in the *zip1* single mutant, although meiosis I promoting factors (i.e., Ndt80, Cdc28, Cdc5, Clb1) remain largely inhibited. These observations imply that the disappearance of the initial signal stimulating the checkpoint is not impacted by *htz1,* placing H2A.Z function downstream in the pathway.

### Influence of H2A.Z on Ndt80 and CDK activity

In our molecular analysis of the *zip1*-induced MRC pathway at various levels, the main alterations detected resulting from the absence of H2A.Z were the dramatic reduction in Ndt80 levels and the persistence of both the Swe1 kinase and phosphorylation of its substrate Cdc18-Y19. The observation that *NDT80* overexpression partially suppresses the *zip1 htz1* arrest raises the possibility that H2A.Z could be directly or indirectly controlling *NDT80* gene expression. It has been recently described that Bdf1, a subunit of the SWR1 complex involved in the interaction with certain histone marks at particular nucleosomes (Altaf *et al.* 2010), is required for meiotic progression and sporulation. Bdf1 binds to the *NDT80* promoter through the BD1 and BD2 bromodomains promoting its transcription (Garcia-Oliver *et al.* 2017). Nevertheless, several observations suggest that H2A.Z does not control Ndt80 levels via Bdf1. The interaction of Bdf1 with the *NDT80* promoter is independent of the SWR1 complex (Garcia-Oliver *et al.* 2017), consistent with our observation that meiotic progression is not significantly affected in the *swr1* single mutant (Figure S2A-S2B). However, the meiotic checkpoint function of H2A.Z does depend on SWR1 since both *zip1 htz1* and *zip1 swr1* show meiotic arrest (Figure S4D). In addition, strong *BDF1* overexpression does not promote sporulation in *zip1 htz1* (Figure S8A). Moreover, we did not find a significant change in *NDT80* transcript levels in our genome-wide expression analysis of the *htz1* mutant during meiosis. Regulation of *NDT80* expression is quite complex and also involves the elimination of the Sum1 repressor binding to the middle-sporulation elements (MSE) in its promoter. The displacement of Sum1 from the MSE requires the competition with Ndt80 and also the phosphorylation of Sum1 by Ime2 and CDK (Winter 2012). We found that, like *zip1 htz1*, the *zip1 htz1 sum1* triple mutant remains blocked in meiosis (Figure S8B) indicating that H2A.Z does not exert its effect on Ndt80 levels via Sum1. In addition, activation of Ndt80 requires its phosphorylation in the nucleus; stimulation of the MRC results in cytoplasmic sequestration of Ndt80 (Wang *et al.* 2011). It is tempting to speculate that H2A.Z could be involved, directly or indirectly, in the nuclear import of Ndt80 when the signal stimulating the checkpoint by the absence of Zip1 declines. The contribution of H2A.Z to the nuclear transport of other proteins has been reported in yeast (Gardner *et al.* 2011), but the almost undetectable levels of Ndt80 in *zip1 htz1* complicate this analysis with the tools currently available.

Our results also show that, in *zip1 htz1*, Swe1-dependent inhibitory phosphorylation of Cdc28-Y19 persists longer than in *zip1*, suggesting that H2A.Z action may be impinging on CDK activity. In fact, deletion of *SWE1*, which abolishes Cdc29-Y19 phosphorylation, significantly suppresses *zip1 htz1* arrest. Since *MIH1*, the gene encoding the phosphatase that reverts Cdc28-Y19 phosphorylation, was found among the genes whose meiotic expression decreases in the *htz1* mutant (Table 1), it is plausible to postulate that lower levels of the Mih1 phosphatase in *zip1 htz1* could explain the accumulation of phosphorylated Cdc28-Y19 and the impaired meiotic progression. However, we demonstrate that strong overproduction of Mih1, which results in negligible Cdc28-Y19 levels, does not restore meiotic nuclear divisions in *zip1 htz1*. This observation, together with the fact that a non-phosphorylatable *cdc28-AF* mutant also has a minimal impact on the kinetics of meiotic progression *of zip1 htz1*, strongly suggest that Swe1 must possess another target in addition to CDK to restrain meiosis in *zip1 htz1*.

Besides CDK, only a limited number of substrates for Swe1/Wee1 have been described. One attractive candidate is Y40 of histone H2B, which is phosphorylated by Swe1 in yeast (or H2B-Y37 phosphorylated by Wee1 in mammals) to control transcription of histone genes (Mahajan *et al.* 2012). H2A.Z interacts with H2B in the nucleosomes; therefore, it is formally possible that the conformational change induced by SWR1-dependent substitution of histone H2A by H2A.Z could modulate the phosphorylation of H2B-Y40 by Swe1. To explore if this chromatin modification has an impact on the MRC, we have generated and analyzed a non-phosphorylatable *htb1-Y40F* mutant and found that the *zip1 htz1 htb2Δ htb1-Y40F* mutant displays the same meiotic arrest as *zip1 htz1* (Figure S8C), indicating that this additional Swe1 target is not relevant for the checkpoint response.

It is surprising that in the *zip1 htz1 cdc28-AF* mutant we observe the induction and accumulation of the proteins involved in meiosis I entry, such as Ndt80, Clb1 and Cdc5, but most cells remain uninucleated (Figure 7). This situation (i.e., accumulation of Ndt80, Cdc5 and Clb1) is reminiscent of the metaphase I arrest induced by a meiotic-depletion *P_CLB2_-cdc20* mutant (Okaz *et al.* 2012) and suggest that, at least some, *zip1 htz1 cdc28-AF* cells are capable of exiting prophase and may arrest at a later stage, such as the metaphase to anaphase I transition. Remarkably, *CLB1* overexpression in *zip1 htz1 cdc28-AF* allows completion of meiotic divisions to a similar degree as does the *zip1 htz1 swe1* mutant. This observation is consistent with the notion that, in the absence of CDK inhibitory phosphorylation (i.e., *cdc28-AF*), Swe1 negatively controls *CLB1* levels in *zip1 htz1* likely by inhibiting a *CLB1*-promoting factor. We note that overexpression of *CLB1* from a high-copy plasmid not only increases the global amount of Clb1, but also accelerates its production being detected at earlier time points in the meiotic kinetics. Execution of proper prophase to meiosis I transition is under tight temporal control by a number of events including the sequential degradation and accumulation of mitotic and meiotic factors, respectively (Okaz *et al.* 2012). We show that *CLB1* overexpression in *zip1 htz1 cdc28-AF* partially restores the proper scenario for timely execution of meiotic transitions. Clb1 is phosphorylated in a CDK- and Cdc5-dependent manner and it is imported to the nucleus by a mechanism that depends on CDK, but not Cdc5 activity. Although Clb1 nuclear localization is not essential for meiotic nuclear divisions it contributes to efficient meiosis I exit (Tibbles *et al.* 2013). On the other hand, the biological relevance of Clb1 phosphorylation remains to be established, but it correlates with the induction of Cdc5. What is the identity of the *CLB1*-promoting factor negatively controlled by Swe1? We speculate that Swe1 could be acting, directly or indirectly, on Ndt80 to inhibit its activity especially in the absence of H2A.Z. We propose a model in which Swe1 action could impact both CDK and Ndt80 activity to restrain meiotic progression (Figure 8A). A cross-talk between CDK and Ndt80 activation in checkpoint-inducing conditions has been also documented (Acosta *et al.* 2011). This model would explain the following situations. 1) In the *zip1 htz1* mutant overexpressing *NDT80*, exogenous levels of this transcription factor could partially overcome Swe1 inhibitory effect on Ndt80, resulting only in a partial release of the meiotic arrest (Figure 6) because Swe1-dependent Cdc28-Y19 phosphorylation would persist (Figure 8B). 2) In the *zip1 htz1 cdc28-AF*, the inhibition of CDK by Swe1 is released because the phosphorylation target is mutated, but the timing of Clb1 induction is incorrect due the opposite effect of CDK and Swe1 on Ndt80 preventing proper meiotic progression (Figure 8C). 3) In the *zip1 htz1 swe1*, both inhibitions on CDK and Ndt80 disappear sustaining meiotic progression (Figure 8D).

In summary, the detailed analysis of the MRC in the *zip1 htz1* allowed us to discover novel functional interactions between the downstream components of the pathway driving meiotic cell cycle progression. Why these aspects are particularly manifested in the absence of H2A.Z? We show here that a number of genes involved in different cell cycle events are misregulated in the *htz1* mutants. A feasible explanation is that the unbalanced levels in cell cycle regulators creates more stringent conditions in *zip1 htz1* for meiosis I entry in comparison with *zip1*, thus revealing more subtle aspects of the molecular mechanisms regulating the prophase to meiosis I transition when the MRC has been deactivated. Additional work will be required to pinpoint the relevant factors targeted by H2A.Z.

**Figure 8.**
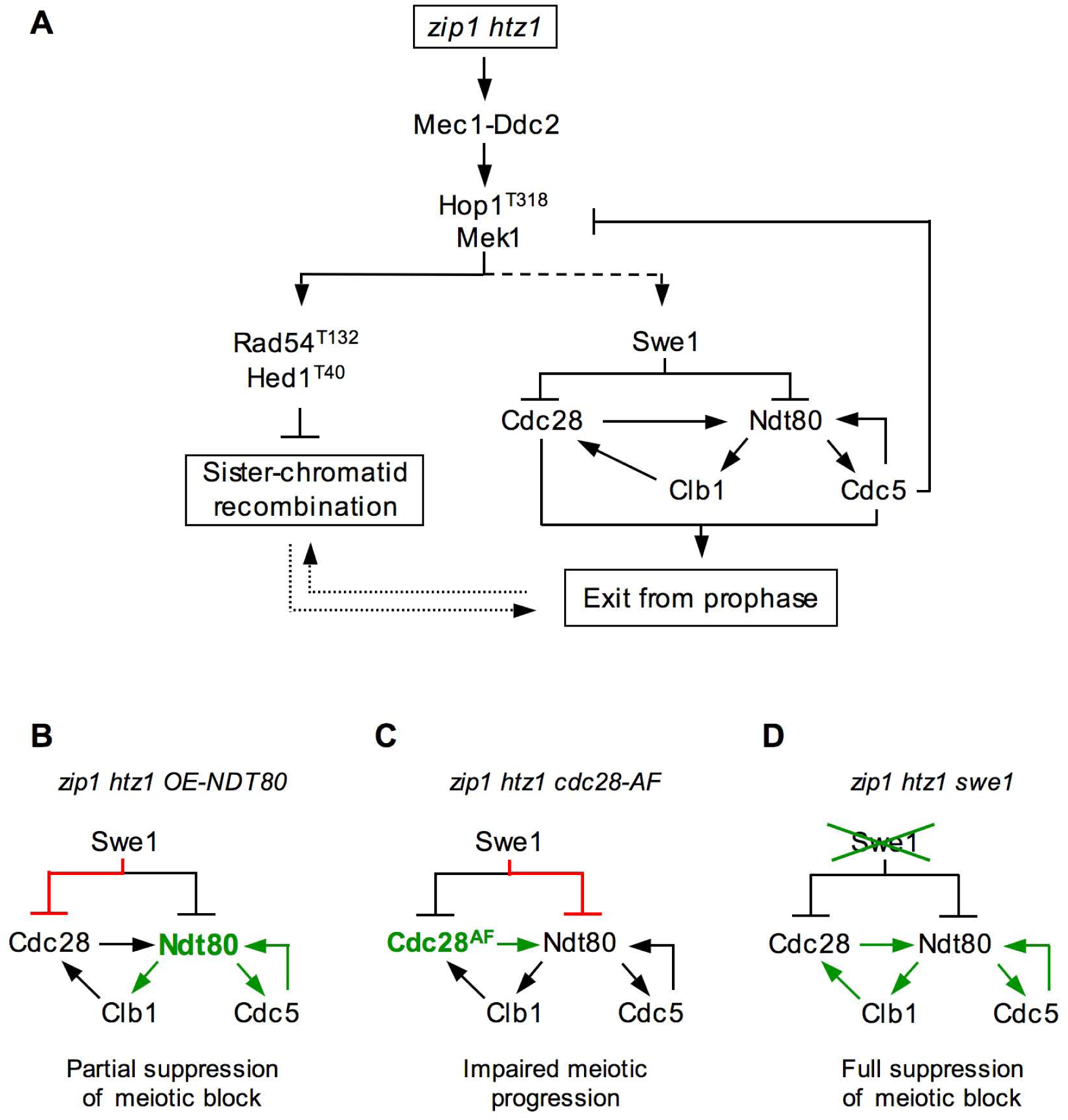
Exit from prophase I in *S. cerevisiae*. (A) A model for the regulation of the prophase to meiosis I transition by the meiotic recombination checkpoint. See text for details. The discontinuous line connecting Mek1 and Swe1 indicates that there is no evidence for direct phosphorylation of Swe1 by Mek1. A functional connection or dependency between DSB repair by sister chromatid recombination and entry into meiosis I is represented by dotted lines. (B), (C) and (D) Impact on meiotic progression resulting from the mutant conditions indicated. Green and red colors represent the predominant positive and negative effects, respectively.

## Acknowledgements

We thank Shirleen Roeder, Jacqueline Segall, Nancy Hollingsworth, Scott Keeney, Michael Lichten, Amy Macqueen, Jesús Carballo, Raimundo Freire and Marisol Santisteban for providing plasmids, strains and/or antibodies. We are also grateful to Irene Gil-Torres and David Núñez for help in strain construction and analysis, to Isabel Acosta for technical assistance and to Andrés Clemente-Blanco and José Pérez-Martín for valuable discussion and suggestions. This work was funded by grants BFU2015-63698-P (to FP), BFU2015-65417-R (to PSS), and BFU2015-69142-REDT (to FP and PSS) from the Ministry of Economy and Competitiveness of Spain (MINECO), and grant CSI084U16 from Junta de Castilla y León (Spain), to PSS.

## SUPPLEMENTAL DATA

**Figure S1.**
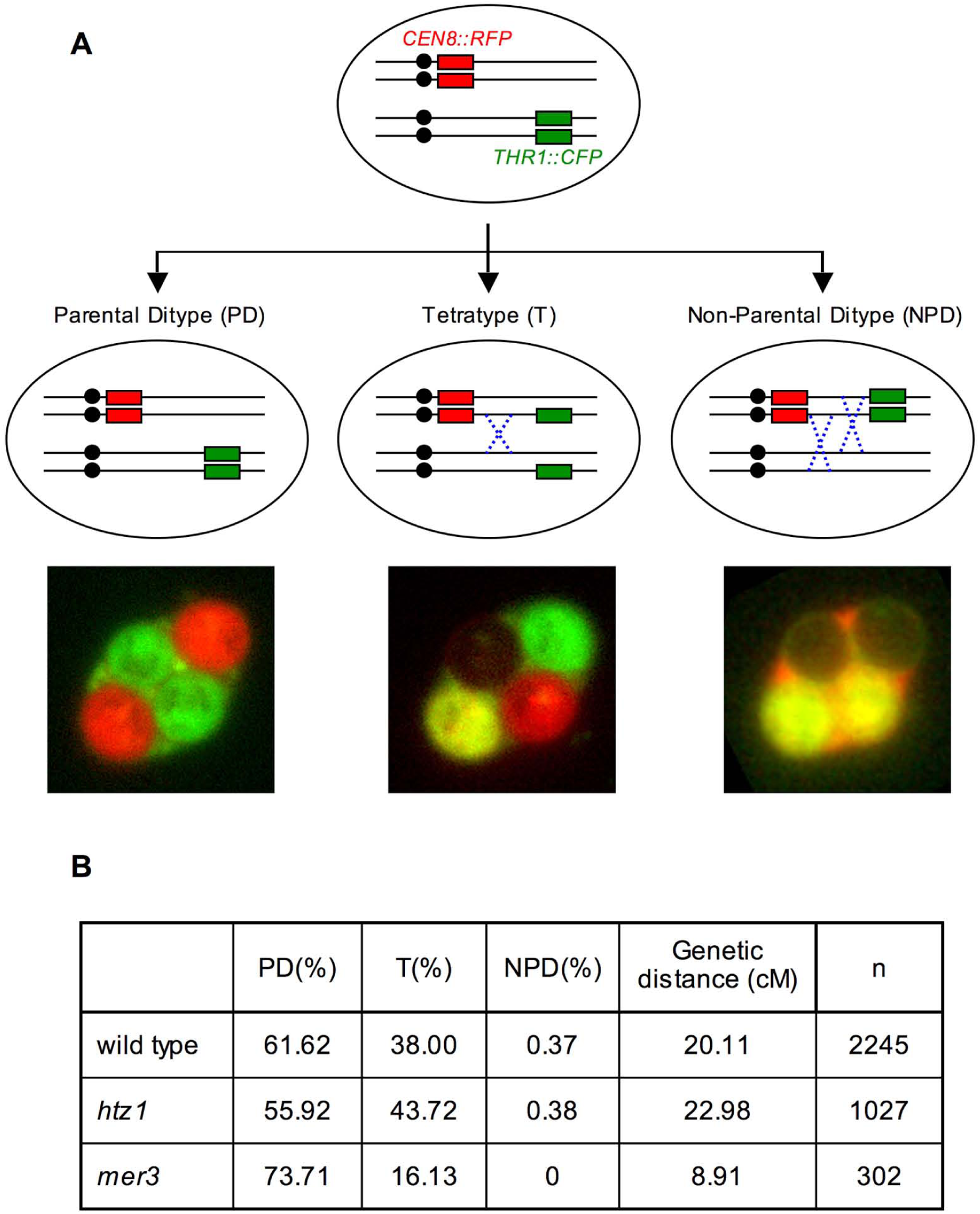
Analysis of crossover frequency by spore-autonomous fluorescence assay. (A): Cartoon depicting the potential configuration of the fluorescent reporter markers on Chromosome VIII and representative images of the different types of asci, according to (Thacker *et al.* 2011). Dashed crosses represent possible recombination events leading to each configuration. (B) The table shows the frequency of PD, T and NPD tetrads, map distance expressed in centiMorgan (cM) and the number of tetrads scored (n) pooled from two inpendent experiments. Strains are DP969 (wild type), DP973 (*htz1*) and DP974 *(mer3).*

**Figure S2.**
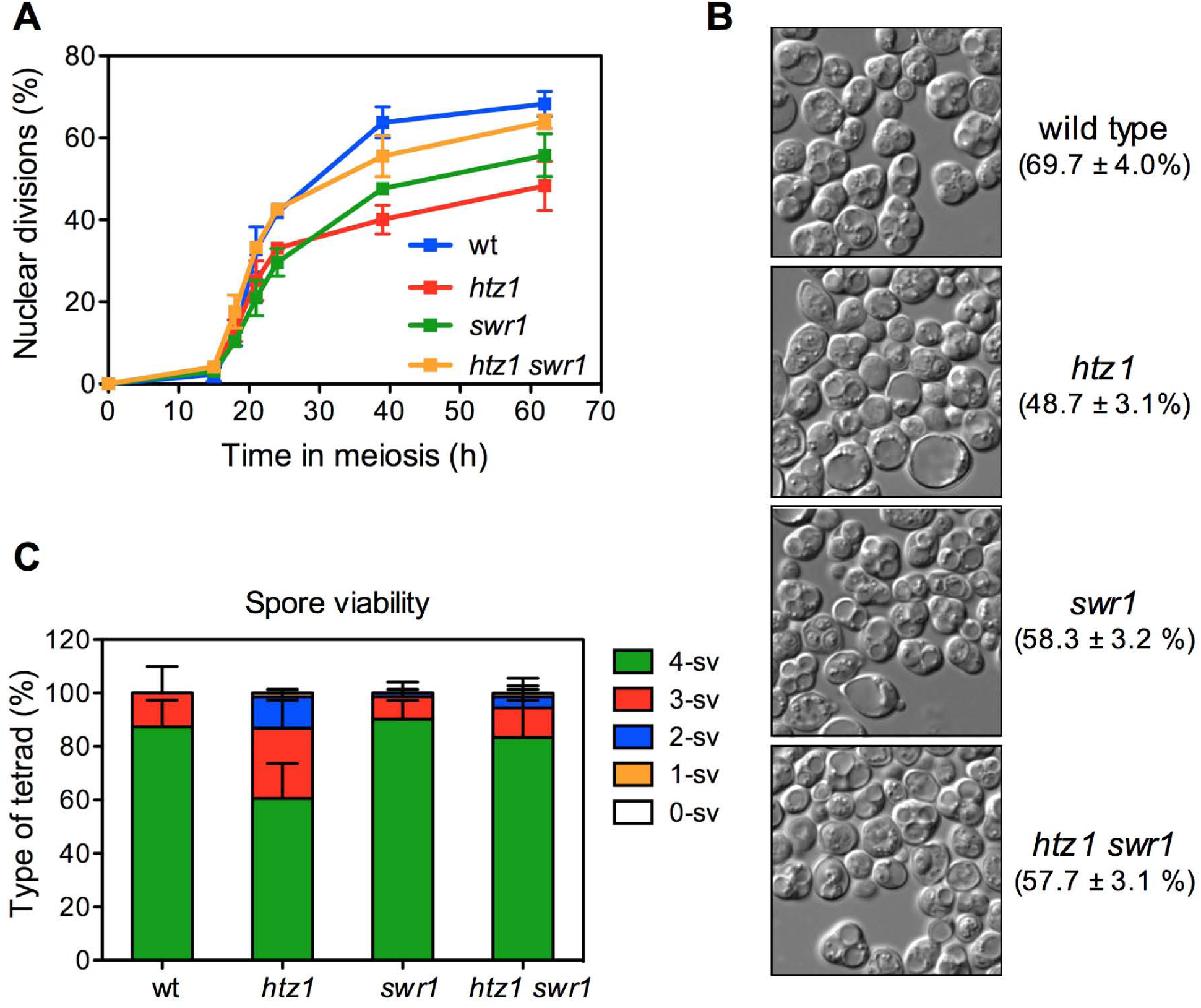
Meiotic impact of the SWRl complex in the presence or absence of H2A.Z. (A) Time course analysis of meiotic nuclear divisions; the percentage of cells containing two or more nuclei is represented. Error bars: range; n=2. (B) Representative DIC images of asci. The sporulation frequency after 62 hours in liquid sporulation rnediurn is shown in parantheses. (C) Spore viability determined by tetrad dissection. The distribution of tetrad types as the percentage of tetrads with 4, 3, 2, 1 and 0 viable spores (4-sv, 3-sv, 2-sv, 1-sv and 0-sv, respectively) is represented. At least 288 spores were scored for each strain Error bars: range n=2. Strains are DP421 (wild type), DP630 (*htz1*), DPI 174 (*swr1*) and DP1056 (*htz1 1 swr1*).

**Figure S3.**
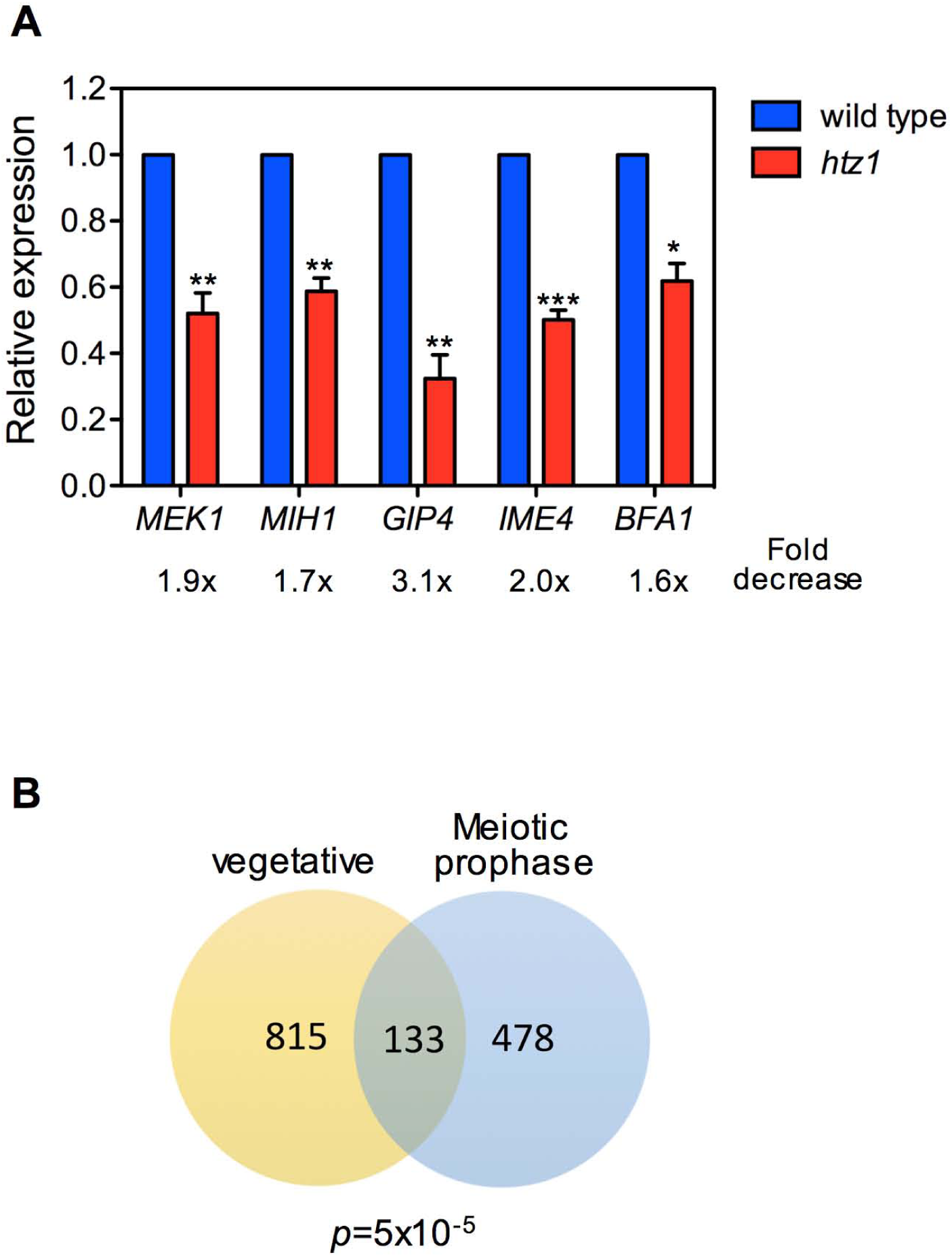
Altered meiotic gene expression in the *htz1* mutant. (A) RT-PCR analysis of mRNA levels of the indicated genes at 15 hours in meiosis. The graph shows relative levels in *htz1* normalized to those in the wild type. Error bars: SD; n=3 (except for *BFA1*; n=2). (B) Venn diagram showing the number of overlapping genes misregulated by *htz1* (1.5-fold cutoff) in vegetative and meiotic cells (15 h). The data for vegetative cells was obtained from (Morillo-Huesca *et al.* 2010). The *p* value was calculated by a hypergeometric test. Strains are DP421 ( wild type) and DP1016 (*htz1*).

**Figure S4.**
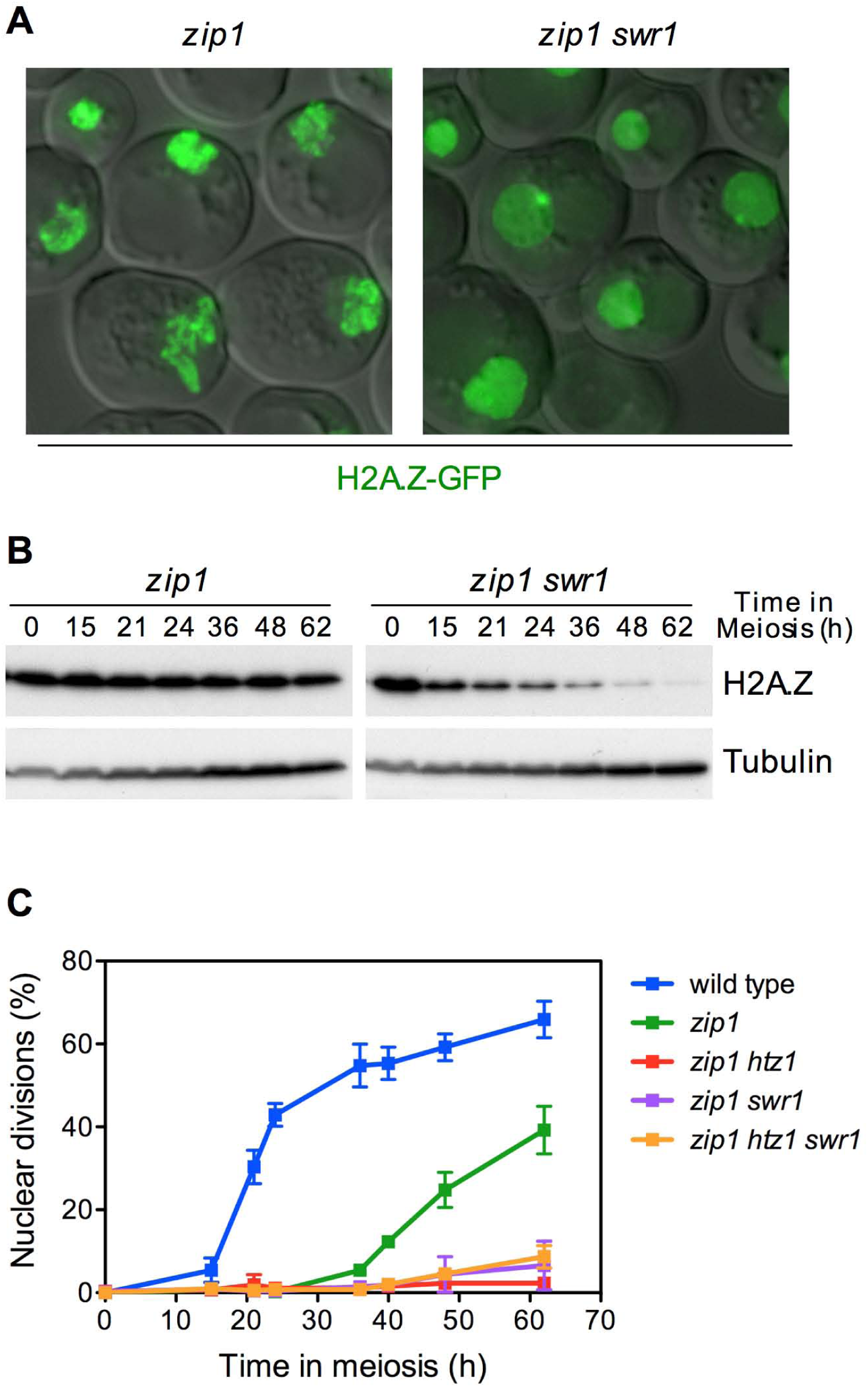
SWR1-dependent chromatin incorporation of H2A.Z is required for normal *zip1*-induced meiotic checkpoint response. (A) Representative images of *zip1* and *zip1 swr1* cells expressing *HTZJ-GFP* at 15 hours after meiotic induction. (B) Western blot analysis of .H2A.Z production during meiosis detected with anti-GFP antibodies. Tubulin was used as a loading control. Strains in (A-B) are DP839 (*zip1 HTZJ-GFP*) and DP842 (*zip1 swr1 HTZ1-GFP*). (C) Time course analysis of meiotic nuclear divisions; the percentage of cells containing two or more nuclei is represented. Error bars: SD; n=3. Strains are DP421 (wild type), DP422 (*zip1*), DP776 (*zip1 htz1*), DP804 (*zip1 swr1*) and DP777 (*zip1 swr1 htz1*).

**Figure S5.**
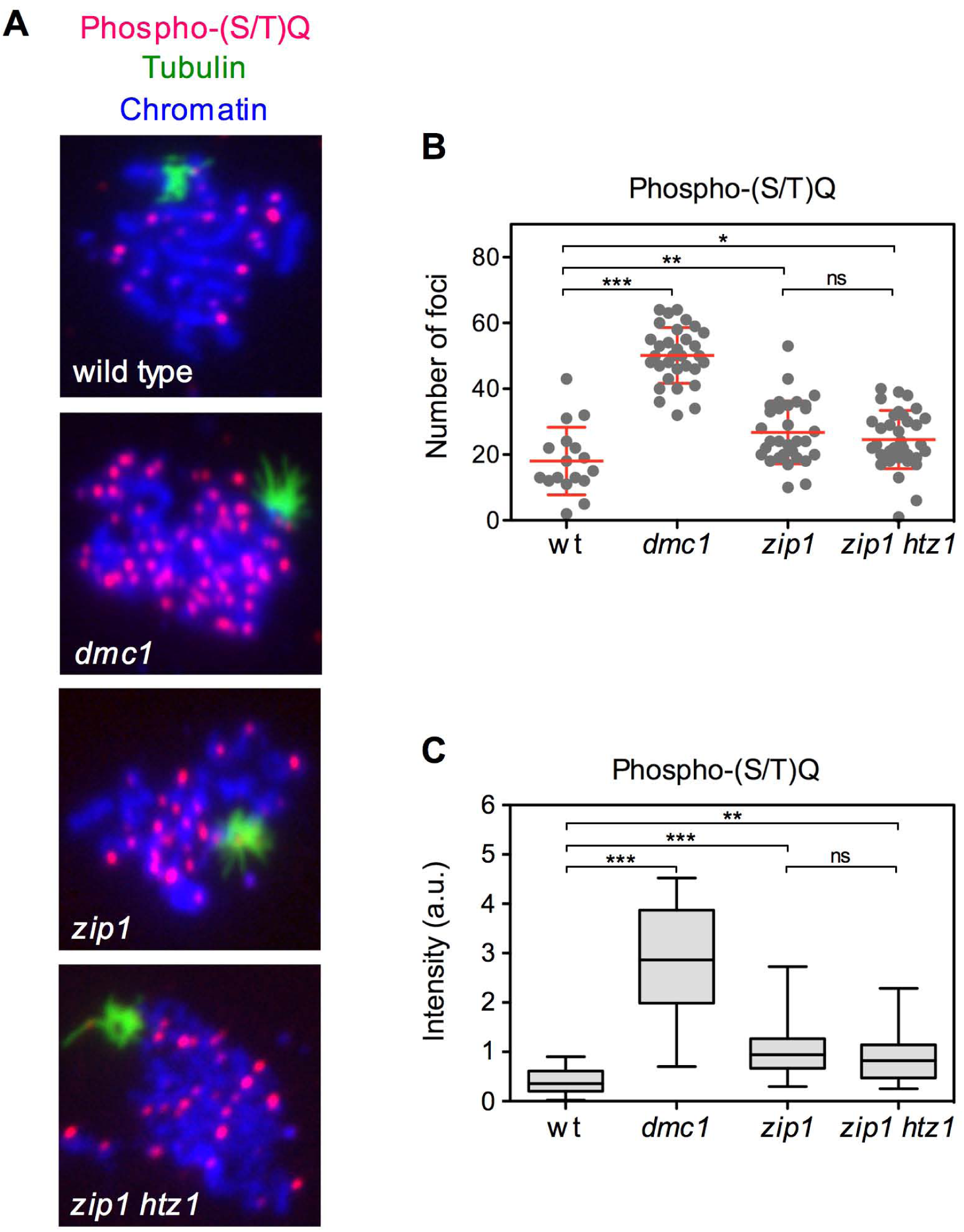
The *zip1 htz1* mutant does not sustain additional Mecl/Tell-dependent DNA damage signaling compared to *zip1.* (A) Immunofluorescence of prophase meiotic chromosomes stained with DAPI (blue), and anti-phospho-(S/T)-Q (red) and anti-tubulin (green) antibodies. The *dmc1* mutant was included as a positive control for the accumulation of extensive meiotic DNA damage (unrepaired resected DSBs). Representative nuclei are shown. (B) Quantification of the number of phospho-(S/T)-Q foci per nucleus. Error bars: SD. (C) Quantification of the fluorescence intensity of the phospho-(S/T)-Q signal per nucleus. Whiskers in the box plot represent the maximum to minimum values. The strains used and the number of nuclei scored in (B-C) are: DP421 (wild type; n=17), DP590 (*dmc1;* n=32), DP1525 (*zip1;* n=31) and DP1526 (*zip1 htz1;* n=34).

**Figure S6.**
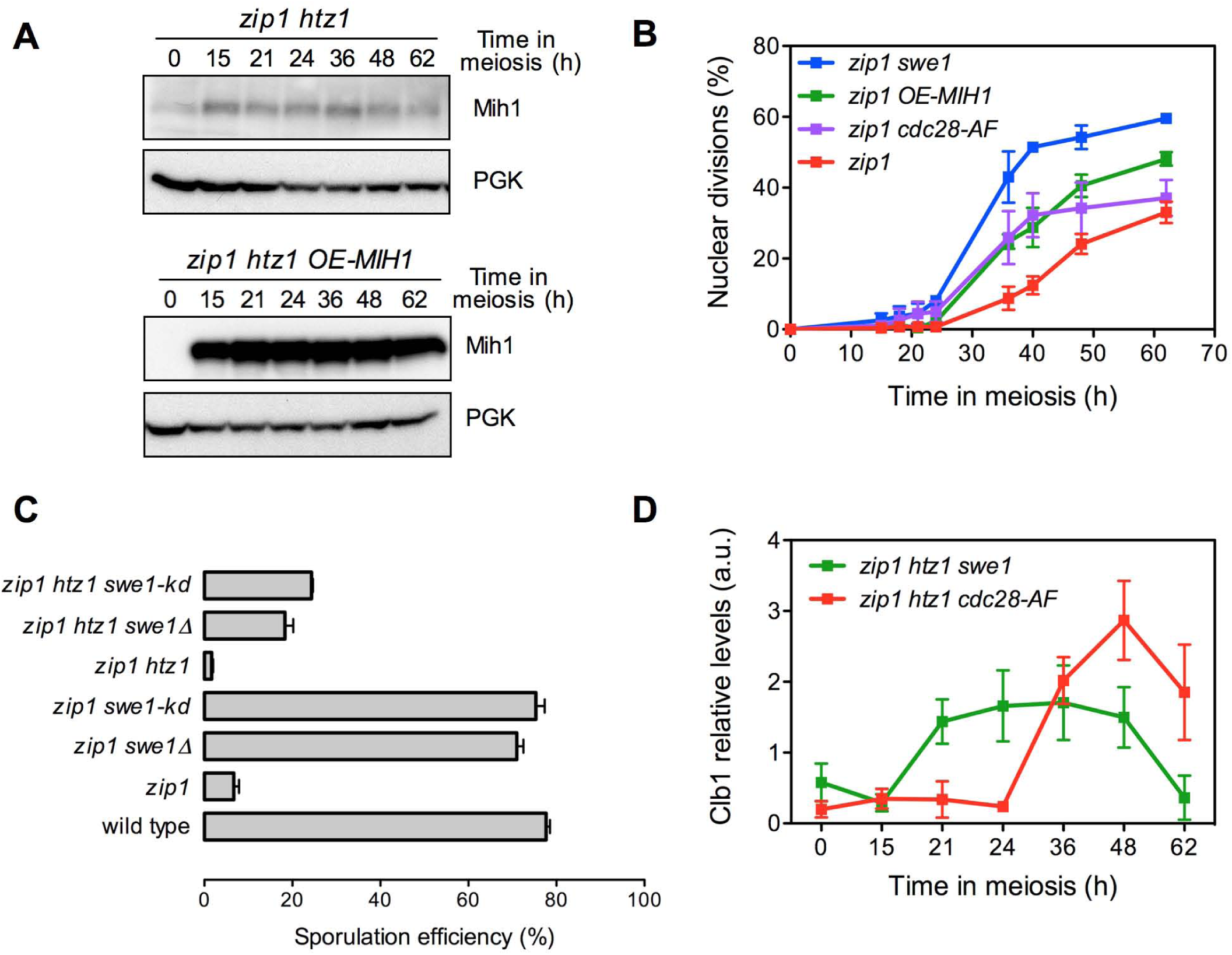
Additional analysis of the role of CDK inhibitory phosphorylation in the iotic checkpoint. (A) Western blot showing *MIH1-GFP* overexpression from the *HOP1* promoter in a high-copy plasmid. The DP1134 strain (*zip1 htz1 MIH1-GFP*) was transformed with vector alone (left panels) or with the pSS265 plasmid (right panels). PGK was used as a loading control. Mih1 was detected with anti-GFP antibodies. (B) Faster meiotic progression in *zip1 cdc28-AF* and *zip1 OE-MIH1* compared to *zip1.* Time course analysis of meiotic nuclear divisions; the percentage of cells containing two or more nuclei is represented. Strains are DP1157 (*zip1 swe1*), DP1153 (*zip1 cdc28-AF*) and DP422 transformed with empty vector pSS248 (*zip1*) or with pSS265 (*zip1 OE-MIH1*). Error bars: SD; n=3. (C) The kinase-dead *swe1-N584A* mutant (*swe1-kd*) phenocopies *SWE1* deletion. The sequence changes introduced to generate the *swe1-N584A* mutation in the genomic locus by *delitto perfetto* are shown. The graph represents the sporulation efficiency determined by microscopic counting as the percentage of cells forming mature or immature asci after 3 days on sporulation plates. Error bars SD; n=3. Strains are: DP1353 (wild type), DP1354 (*zip1*), DP1157 (*zip1 swe1*) and DP1467 (*zip1 swe1-kd*), DP1414 (*zip1 htz1*), DP1113 *( zipl htz1 swel)* and DP1468 (*zip1 htz1 1-kd*). (D) Quantification of Clb1 levels, normalized to PGK, throughout meiosis in the indicated strains. Error bars: SD; n=3. Strains are DP1113 (*zip1 htz1 swel*) and DP1416 (*zip1 htz1 cdc28-AF).*

**Figure S7.**
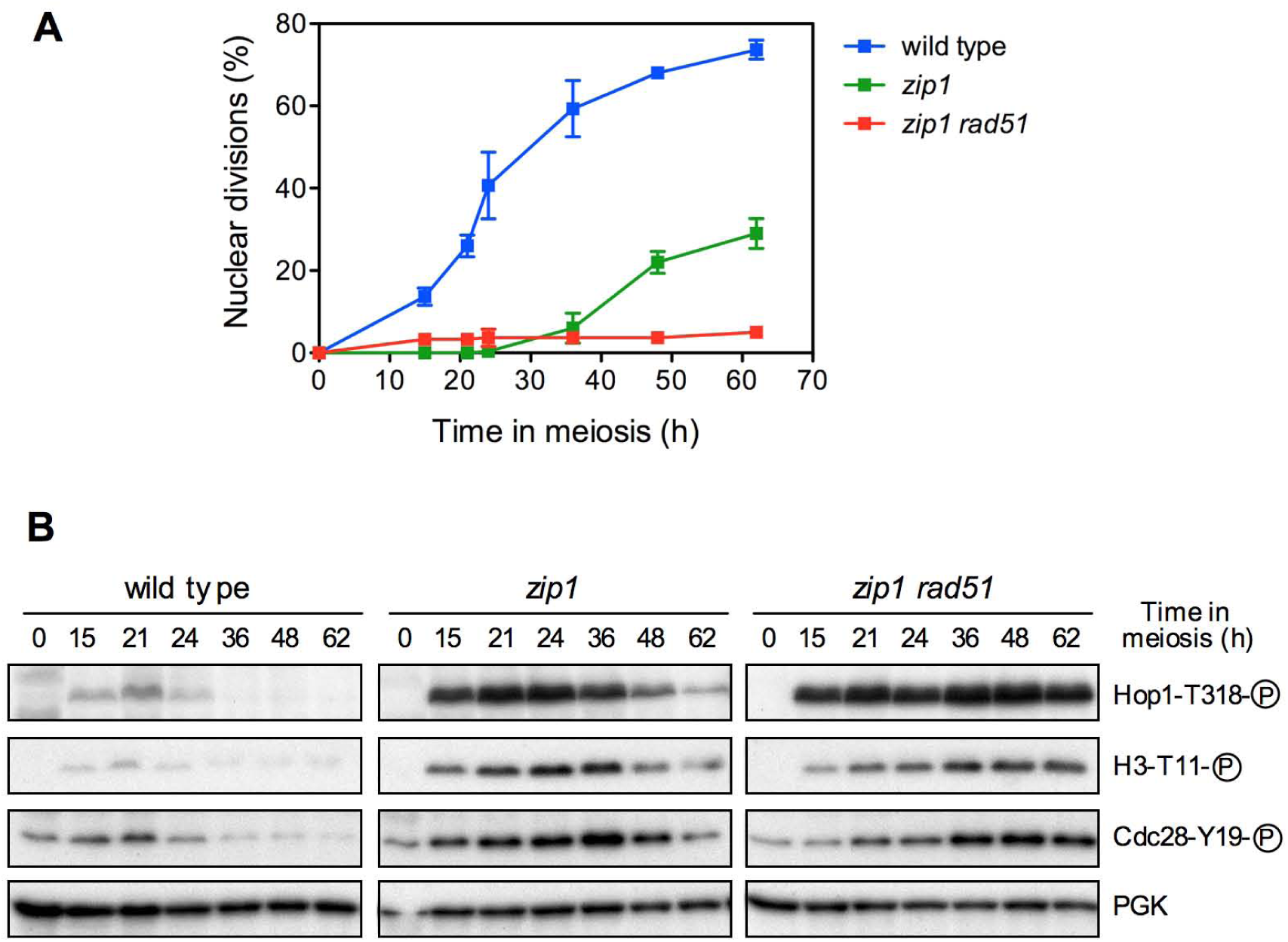
Deletion of *RAD51* leads to sustained checkpoint activation and meiotic arrest *zip1.* (A) Time course analysis of meiotic nuclear divisions; the percentage of cells containing two or more nuclei is represented. Error bars: SD; n=3. (B) Western blot analysis of the indicated molecular markers of checkpoint activity. PGK was used as a loading control. strains are DP1359 (wild type), DP1360 (*zip1*) and DP1364 (*zip1 rad5 1*)

**Figure S8.**
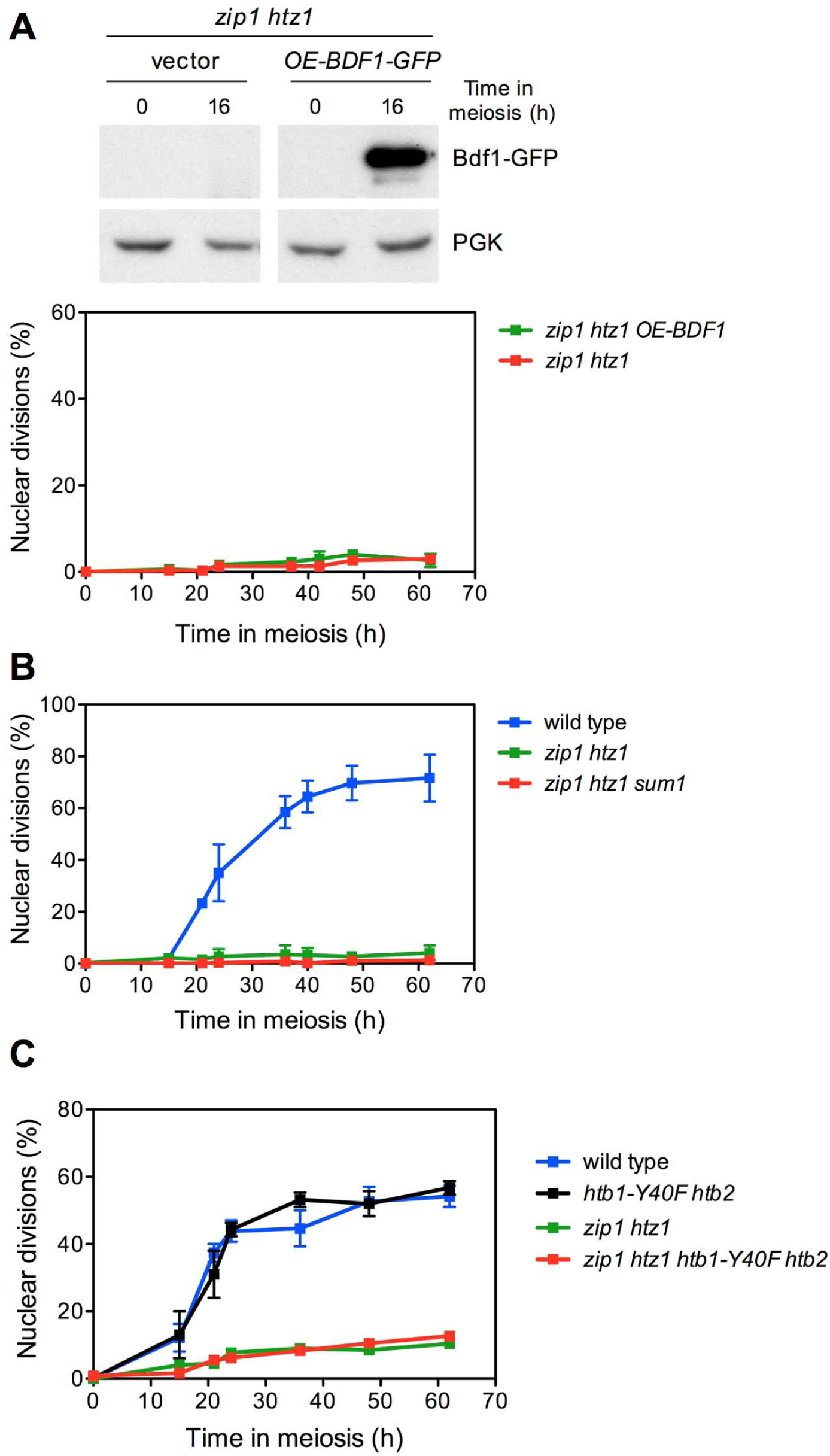
Meiosis-specific overexpression of *BDF1,* deletion of *SUM 1* and mutation of *H2B-Y40* do not suppress *zip1 htz1* arrest. (A) Upper panel; western blot analysis of *zip1 htz1* cells (DP1017) transformed with vector (pSS248) or with a 2-micron plasmid expressing *BDF1-GFP* from the *HOPI* promoter (*OE-BDF1-GFP;* pSS354). Extracts were prepared at time zero or 16 hours after meiotic induction (around the peak of prophase) and analyzed with anti-GFP antibodies. PGK was used as loading control. Lower panel; time course analysis of meiotic nuclear divisions in the ame strains. The percentage of cells containing two or more nuclei is represented. Error bars: SD; n=3. (B) Time course analysis of meiotic nuclear divisions in the indicated strains. Wild type (DP421), *zip1 htz1* (DP630) and *zip1 htz1 sum1* (DP1441). Error bars: range; n=2. (C). Time course analysis of meiotic nuclear divisions in the indicated strains. Wild type (DP1445), *htb1-Y40F htb2* (DP1446), *zip1 htz1* (DP1449) and *zip1 htz1 htb1-Y40F htb2* (DP1450). Error bars: range; n=2.

**Table S1.** *Saccharomyces cerevisiae* strains.

| Strain | Genotype | Source |
| --- | --- | --- |
| DP421 | *MATa/MATα leu2,3-112 his4-260 thr1-4 ura3-1 trp1-289 ade2-1 lys2ΔNheI* | PSS Lab |
| DP422 | DP421 *zip1::LYS2* | PSS Lab |
| DP437 | DP421 *ZIP1-GFP* | PSS Lab |
| DP449 | DP421 *zip1::LYS2 DDC2-GFP::TRP1* | PSS Lab |
| DP590 | DP421 *dmc1::hphMX4 p306(BrdU-Inc)::URA3/ura3-1* | PSS Lab |
| DP630 | DP421 *htz1::URA3* | This work |
| DP631 | DP421 *zip1::LYS2 htz1::URA3* | This work |
| DP713 | DP421 *mek1::kanMX6* | PSS Lab |
| DP776 | DP421 *zip1::LYS2 htz1::URA3 DDC2-GFP::TRP1* | This work |
| DP777 | DP421 *zip1::LSY2 htz1::URA3 swr1::natMX4 DDC2-GFP::TRP1* | This work |
| DP804 | DP421 *swr1::natMX4 zip1::LSY2 DDC2-GFP::TRP1* | This work |
| DP815 | DP421 *zip1::LYS2 htz1::URA3 spo11::natMX4 DDC2-GFP::TRP1* | This work |
| DP816 | DP421 *zip1::LYS2 htz1::URA3 ddc2::TRP1 sml1::KanMX6* | This work |
| DP838 | DP421 *ZIP1-GFP htz1::URA3* | This work |
| DP839 | DP421 *zip1::LYS2 HTZ1-GFP::kanMX6* | This work |
| DP840 | DP421 *HTZ1-GFP::kanMX6* | This work |
| DP841 | DP421 *swr1::natMX4 HTZ1-GFP::kanMX6* | This work |
| DP842 | DP421 *zip1::LYS2 swr1::natMX4 HTZ1-GFP::kanMX6* | This work |
| DP969 | SK1 *CEN8::tdTomato-LEU2/CEN8 THR1:mCerulean-TRP1/THR1* | S. Keeney |
| DP973 | SK1 *CEN8::tdTomato-LEU2/CEN8 THR1:mCerulean-TRP1/THR1 htz1::hphMX4* | This work |
| DP974 | SK1 *CEN8::tdTomato-LEU2/CEN8 THR1:mCerulean-TRP1/THR1 mer3::hphMX4* | This work |
| DP1016 | DP421 *htz1::hphMX4* | This work |
| DP1017 | DP421 *zip1::LYS2 htz1::hphMX4* | This work |
| DP1056 | DP421 *htz1::URA3 swr1:: natMX4* | This work |
| DP1113 | DP421 *zip1::LYS2 htz1::hphMX4 swe1::natMX4* | This work |
| DP1134 | DP421 *zip1::LYS2 htz1::hphMX4 MIH1-GFP::kanMX6* | This work |
| DP1144 | DP421 *htz1::URA3 spo11::ADE2* | This work |
| DP1153 | DP421 *zip1::LYS2 cdc28-AF::LEU2* | This work |
| DP1154 | DP421 *zip1::LYS2 htz1::hphMX4 cdc28-AF::LEU2* | This work |
| DP1157 | DP421 *zip1::LYS2 swe1::LEU2* | This work |
| DP1174 | DP421 *swr1::hphMX4* | This work |
| DP1185 | DP421 *TPR1-P<sub>GALI</sub>-ZIP1-GFP ura3::P<sub>GPD1</sub>-GAL4(848)ER::URA3/ura3-1* | This work |
| DP1186 | DP421 *TPR1-P<sub>GALI</sub>-ZIP1-GFP P<sub>GPD1</sub>-GAL4(848)ER::URA3/ura3-1 htz1::hphMX4* | This work |
| DP1259 | DP421 *htz1::URA3 mek1::kanMX6* | This work |
| DP1353 | DP421 *3MYC-SWE1* | This work |
| DP1354 | DP421 *zip1::LYS2 3MYC-SWE1* | This work |
| DP1359 | DP421 *TUB1/TUB1-GFP::TRP1* | PSS Lab |
| DP1360 | DP421 *zip1::LYS2 TUB1/TUB1-GFP::TRP1* | PSS Lab |
| DP1364 | DP421 *zip1::LYS2 rad51::natMX4 TUB1/TUB1-GFP::TRP1* | PSS Lab |
| DP1414 | DP421 *zip1::LYS2 htz1::hphMX4 3MYC-SWE1* | This work |
| DP1416 | DP421 *zip1::LYS2 htz1::hphMX4 cdc28-AF::LEU2 3MYC-SWE1* | This work |
| DP1441 | DP421 *zip1::LYS2 htz1::URA3 sum1::natMX4* | This work |
| DP1445 | DP421 *[hta1-htb1]::kanMX6 [hta2-htb2]::natMX4*<br>pSS347 (*HTA1-HTB1*)-*TRP1* | This work |
| DP1446 | DP421 *[hta1-htb1]::kanMX6 [hta2-htb2]::natMX4*<br>pSS348 (*HTA1-htb1-Y40F*)-*TRP1* | This work |
| DP1449 | DP421 *zip1::LYS2 htz1::hphMX4 [hta1-htb1]::kanMX6 [hta2-htb2]::natMX4*<br>pSS347 (*HTA1-HTB1*)-*TRP1* | This work |
| DP1450 | DP421 *zip1::LYS2 htz1::hphMX4 [hta1-htb1]::kanMX6 [hta2-htb2]::natMX4*<br>pSS348 (*HTA1-htb1-Y40F*)-*TRP1* | This work |
| DP1467 | DP421 *zip1::LYS2 3MYC-swe1-N584A* | This work |
| DP1468 | DP421 *zip1::LYS2 htz1::hphMX4 3MYC-swe1-N584A* | This work |
| DP1523 | DP421 *spo11::ADE2* | This work |
| DP1524 | DP421 *zip1::LYS2 spo11::ADE2* | This work |
| DP1525 | DP421 *zip1::LYS2 p306(BrdU-Inc)::URA3/ura3-1* | This work |
| DP1526 | DP421 *zip1::LYS2 htz1::hphMX4 p306(BrdU-Inc)::URA3/ura3-1* | This work |

\* All strains are diploids isogenic to BR1919 (Rockmill and Roeder 1990), except strains DP969, DP973 and DP974, which are diploids isogenic to SK1. The haploid parents of DP969 (SK1) were obtained from S. Keeney (Thacker *et al.* 2011). Unless specified, all strains are homozygous for the indicated markers. DP421 is a *lys2* version of the original BR1919-2N.
Rockmill, B., and G. S. Roeder, 1990 Meiosis in asynaptic yeast. *Genetics* 126: 563-574.
Thacker, D., I. Lam, M. Knop and S. Keeney, 2011 Exploiting spore-autonomous fluorescent protein expression to quantify meiotic chromosome behaviors in *Saccharomyces cerevisiae*. *Genetics* 189: 423-439.

**Table S2.** Plasmids

| **Plasmid** | **Vector** | **Relevant parts** | **Source/Reference** |
| --- | --- | --- | --- |
| pTK17 | pUC19 | *htz1::URA3* | (SANTISTEBAN *et al.* 2000) |
| pJC29 | pRS426 | *2μ URA3 CDC5* | (JASPERSEN *et al.* 1998) |
| pR2042 | pRS305 | *LEU2 cdc28-AF* | (LEU and ROEDER 1999) |
| pR2045 | pRS426 | *2μ URA3 CLB1* | (LEU and ROEDER 1999) |
| pSS263 | pRS426 | *2μ URA3 NDT80* | This work |
| pSS248 | pYES2 derivative | *2μ URA3 P<sub>HOP1</sub>-GFP* | This work |
| pSS265 | pSS248 | *2μ URA3 P<sub>HOP1</sub>-GFP-MIH1* | This work |
| pSS345 | pRS316 | *CEN6 URA3 HTA1-HTB1* | This work |
| pSS347 | pRS314 | *CEN6 TRP1 HTA1-HTB1* | This work |
| pSS348 | pRS314 | *CEN6 TRP1 HTA1-htb1-Y40F* | This work |
| pSS354 | pSS248 | *2μ URA3 P<sub>HOP1</sub>-GFP-BDF1* | This work |

JASPERSEN, S. L., J. F. CHARLES, R. L. TINKER-KULBERG and D. O. MORGAN, 1998 A late mitotic regulatory network controlling cyclin destruction in *Saccharomyces cerevisiae*. *Mol Biol Cell* **9**: 2803-2817.
LEU, J. Y., and G. S. ROEDER, 1999 The pachytene checkpoint in *S. cerevisiae* depends on Swe1-mediated phosphorylation of the cyclin-dependent kinase Cdc28. *Mol Cell* **4**: 805-814.
SANTISTEBAN, M. S., T. KALASHNIKOVA and M. M. SMITH, 2000 Histone H2A.Z regulates transcription and is partially redundant with nucleosome remodeling complexes. *Cell* **103**: 411-422.

**Table S3.**
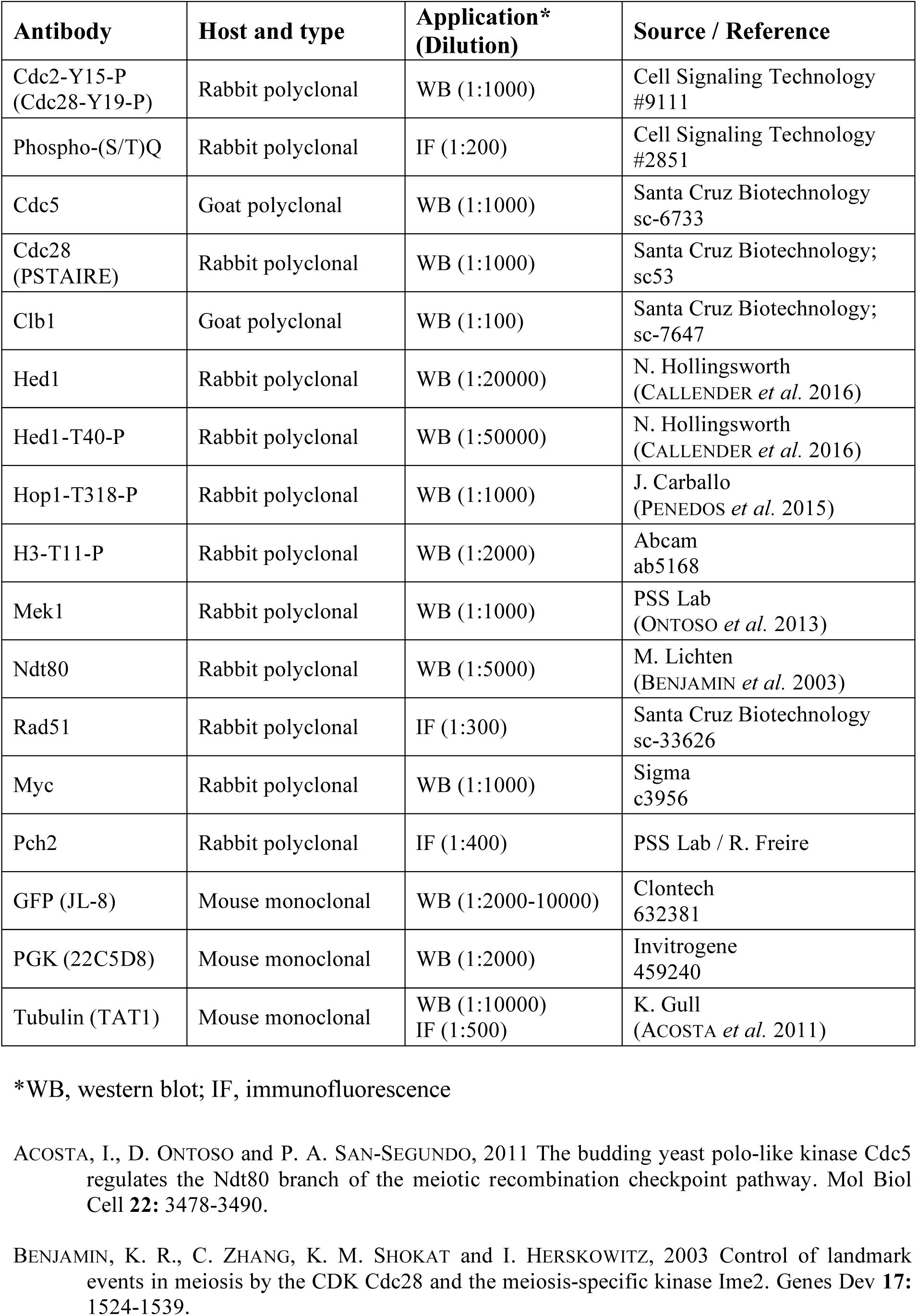
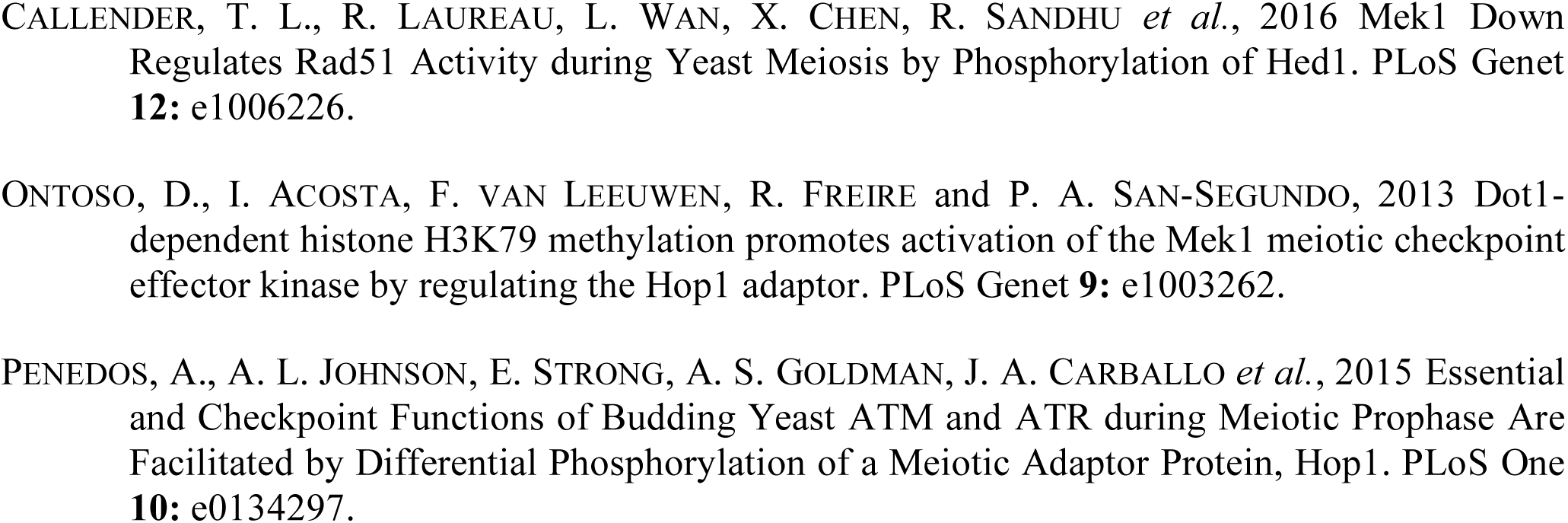
Primary antibodies

